# Substrate dynamics contribute to enzymatic specificity in human and bacterial methionine adenosyltransferases

**DOI:** 10.1101/2021.02.18.431797

**Authors:** Madhuri Gade, Li Lynn Tan, Adam M. Damry, Mahakaran Sandhu, Jospeh S. Brock, Andie Delaney, Alejandro Villar-Briones, Colin J. Jackson, Paola Laurino

**Affiliations:** Protein Engineering and Evolution Unit, Okinawa Institute of Science and Technology Graduate University, 1919-1 Tancha, Onna, Okinawa, Japan 904-0495; Research School of Chemistry, Australian National University, Canberra, 2601, Australia; Research School of Biology, Australian National University, Canberra, 2601, Australia; Australian Research Council Centre of Excellence for Innovations in Peptide and Protein Science, Research School of Chemistry, Australian National University, Canberra, 2601, ACT, Australia; Australian Research Council Centre of Excellence in Synthetic Biology, Research School of Chemistry, Australian National University, Canberra, 2601, ACT, Australia

## Abstract

Protein conformational change can facilitate the binding of non-cognate substrates and underlie promiscuous activities. However, the contribution of substrate conformational dynamics to this process is comparatively poorly understood. Here we analyse human (hMAT2A) and *Escherichia coli* (eMAT) methionine adenosyltransferases that have identical active sites but different substrate specificity. In the promiscuous hMAT2A, non-cognate substrates bind in a stable conformation to allow catalysis. In contrast, non-cognate substrates rarely sample stable productive binding modes in eMAT owing to altered mobility in the enzyme active site. Different cellular concentrations of substrate likely drove the evolutionary divergence of substrate specificity in these orthologs. The observation of catalytic promiscuity in hMAT2A led to the detection of a new human metabolite, methyl thioguanosine, that is produced at elevated level in a cancer cell line. This work establishes that identical active sites can result in different substrate specificity owing to the effects of substrate and enzyme dynamics.

## Introduction

Enzymes can exhibit promiscuous activities with non-cognate substrates that are not involved in the main physiological function of the enzyme^1^. These promiscuous activities are often vestigial traits of a distant ancestor^2^ or have originated by chance through evolution^3–6^. The importance of promiscuous enzymatic activities is becoming increasing evident, as they have been shown to contribute to evolvability^7^, stress responses^8^ and, potentially, susceptibility to disease^8–10^. Protein conformational sampling has been shown to play a role in substrate promiscuity^11–14^, as conformational change can allow enzyme to occasionally sample alternative conformations with different charge preorganization, allowing different transition states to be stabilized^15^. While the role of protein structural dynamics in this process has been described, the role of substrate conformational sampling is comparatively poorly understood. It has recently been reported that large active sites can accommodate multiple different productive substrate conformations without changing the conformation of the catalytic pocket^16, 17^, and that in some cases new Michaelis complexes can be recognized^18^.

The methionine adenosyltransferases (MATs), are found in all kingdoms of life and the product of their reaction, S-adenosyl-L-methionine (SAM), is a necessary metabolite in several essential cellular processes^19–21^. Because of the physiological importance of SAM, dysfunction in the production of SAM by MATs can lead to disease^22,23^. Mechanistically, the enzyme-catalysed formation of S-adenosyl-L-methionine (SAM) from adenosine triphosphate (ATP) and methionine occurs in two steps^24^: first, SAM is formed by S_N_2 attack by the sulfur of methionine at the C5’ carbon of ATP, followed by hydrolysis of triphosphate (PPPi) into pyrophosphate (PPi) and orthophosphate (Pi)^25^ (Figure 1a). This second step is believed to provide the energy required for the conformational rearrangement of the enzyme necessary for product release^26^. Two Mg^2+^ ions are involved in coordination of the triphosphate moiety of ATP and K^+^ is known to enhance the reaction rate by allowing the active site to adopt the optimal conformation^20, 27, 28^.

**Figure 1.**
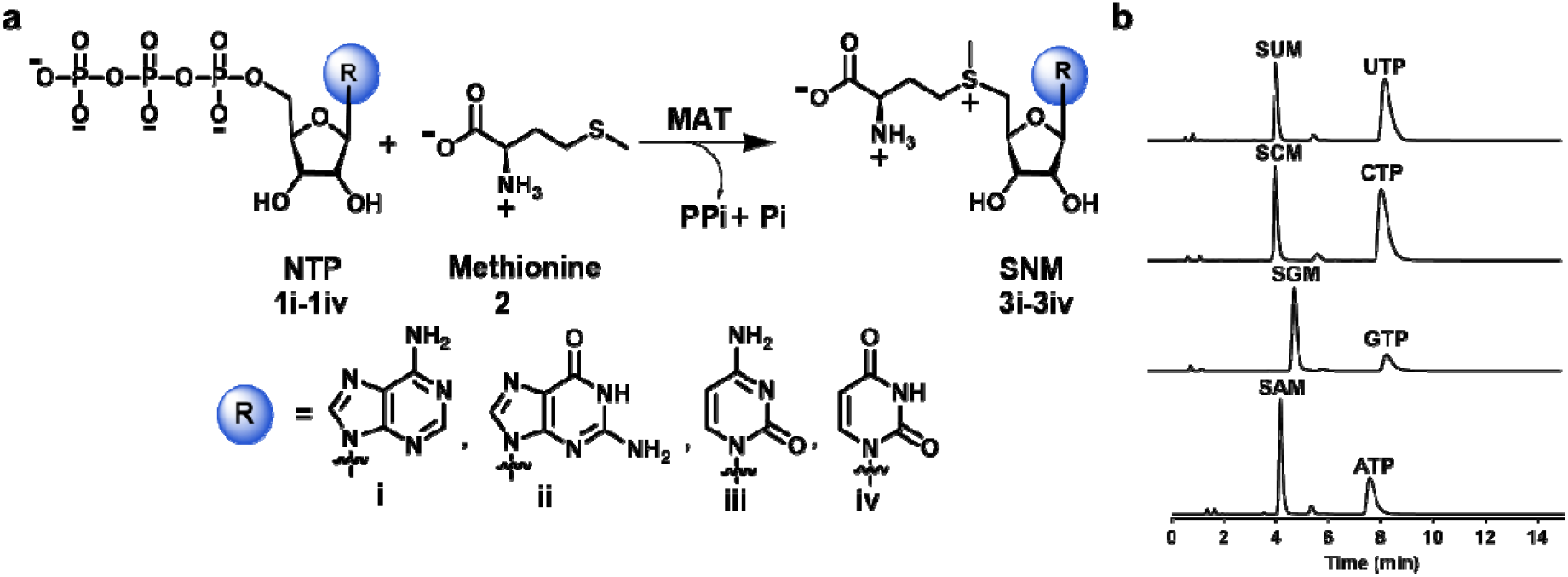
SNM biochemical synthesis, identification of SNM analogs by UPLC. **a)** Synthesis of S-nucleoside-L-methionine (SNM) analogs S-adenosyl-L-methionine (3i, SAM), S-guanosyl-L-methionine (3ii, SGM), S-cytidyl-L-methionine (3iii, SCM), S-uridyl-L-methionine (3iv, SUM) from different nucleotides (ATP, GTP, CTP, UTP) and methionine. **b)** UPLC chromatograms of the reaction of NTPs (5 mM) and methionine (10 mM) in presence of hMAT2A (20 μM) (1h, 37 °C, details are in the Methods section). Noted are the peaks corresponding to SAM (t_R_ = 4.1 min), SCM (t_R_ = 4.6 min), SUM (t_R_ = 4.6 min), SGM (t_R_ = 5.3 min), ATP (t_R_ = 7.5 min), GTP (t_R_ = 7.8 min), CTP (t_R_ = 8.3 min) and UTP (t_R_ = 8.5 min).

MATs are an excellent model system for the study of substrate promiscuity because the chemical reactivity of the cognate physiological nucleotide substrate, ATP, is independent from the nucleobase. The C5’ atom, which acts as electrophile in the MAT-catalysed reaction, belongs to the sugar moiety of the nucleotide, and is therefore distant from the nucleobase^29, 30^. Moreover, SAM is not an intrinsically better methyl donor than the potential products from promiscuous reactions with non-cognate NTPs (S-guanosyl-L-methionine (SGM), S-cytidyl-L-methionine (SCM), or S-uridyl-L-methionine (SUM)), since the nucleobase does not influence the sulfonium reactivity. While *E. coli* MAT has been reported to display specificity for ATP *in vitro*^27^, the promiscuity of human MAT for GTP, CTP and UTP has not been systematically explored *in vitro* or *in vivo*.

In this work, we have performed a systematic study of the substrate promiscuities of human (hMAT2A) and *E. coli* (eMAT) MATs. We show that hMAT2A, unlike eMAT, exhibits substrate promiscuity towards other non-cognate NTPs. Structural analysis reveals that eMAT specificity is a consequence of altered structural constraints on non-cognate substrates in combination with increased active site loop dynamics *vs*. hMAT2A. The increased conformational freedom of the non-cognate substrates results in eMAT sampling catalytically non-productive states at higher frequency than the native substrate, ATP, providing a molecular explanation for the observed enzyme kinetics. We demonstrate that the substrate promiscuity of hMAT2A is relevant *in vivo*, and this knowledge allowed us to identify a new metabolite, methyl thioguanosine, a breakdown product of SGM, that is produced in a human liver cancer cell line but was not produced at detectable levels in a normal liver cell line.

## Results

### The catalytic promiscuity of MATs

For the kinetic analysis of eMAT and hMAT2A, we developed a sensitive and specific assay based on ultra-performance liquid chromatography (UPLC) (see Methods). This allowed us to analyse the generation of different S-(nucleoside)-L-methionine (SNM) analogs with confirmation of the products *via* mass spectrometry (Figure 1b; Appendix). From these data, kinetic parameters were derived (Table 1; Supplementary Figure 1). hMAT2A efficiently catalysed the formation of SGM, SCM and SUM, in addition to the cognate product, SAM. No spontaneous product formation was observed without MAT (Supplementary Figure 2a). The activity of hMAT2A with the four different nucleotides varied over a relatively narrow range (*k*_cat_/*K*_M_ values of non-cognate substrates within 42-93% of ATP; Table 1). The *k*_cat_/*K*_M_ of eMAT with ATP was comparable to that of hMAT2A (Table 1), albeit with lower *k*_cat_ and *K*_M_ values. However, the activity of eMAT for other nucleotides differed: the *k*_cat_/*K*_M_ of eMAT was 61-fold, 8.5-fold, and 139-fold lower for GTP, CTP and UTP respectively, in comparison to ATP (Table 1; Supplementary Figure 1). Thus, while hMAT2A is catalytically promiscuous with various NTPs, eMAT is comparatively specific (Supplementary Figure 2b).

**Table 1.** Kinetic parameters* for the SNM analog formation by hMAT2A and eMAT.

| Enzyme:Substrate | $k_{cat}$ (s<sup>-1</sup>) | $K_M$ (mM) | $k_{cat}/K_M$ (M<sup>-1</sup>s<sup>-1</sup>) |
| --- | --- | --- | --- |
| hMAT2A:ATP | 764 ± 61 | 0.27 ± 0.07 | 2.8 × 10<sup>6</sup> |
| hMAT2A:GTP | 3270 ± 600 | 1.26 ± 0.40 | 2.6 × 10<sup>6</sup> |
| hMAT2A:CTP | 100 ± 4.8 | 0.08 ± 0.02 | 1.3 × 10<sup>6</sup> |
| hMAT2A:UTP | 1180 ± 140 | 0.97 ± 0.2 | 1.2 × 10<sup>6</sup> |
| eMAT:ATP | 66 ± 4 | 0.06 ± 0.02 | 1.1 × 10<sup>6</sup> |
| eMAT:GTP | 18 ± 2 | 0.97 ± 0.30 | 1.8 × 10<sup>4</sup> |
| eMAT:CTP | 168 ± 12 | 1.3 ± 0.25 | 1.3 × 10<sup>5</sup> |
| eMAT:UTP | 23 ± 3 | 2.90 ± 0.7 | 7.9 × 10<sup>3</sup> |

\* Kinetic parameters for the SNM analog formation by hMAT2A, eMAT using a concentration of ATP, GTP, CTP and UTP in the range of 0.025 to 5 mM and a fixed saturating concentration of methionine (10 mM) in the presence of HEPES (100 mM), KCl (50 mM), MgCl<sub>2</sub> (10 mM), pH 8 at 37 °C. [hMAT2A] was 0.5 μM and [eMAT] concentrations were 0.5 μM for ATP, 5 μM for GTP, CTP and 10 μM for UTP. SNM production was analyzed by UPLC, and data fitted to the Michaelis-Menten equation using GraphPad Prism 7.02 (Supplementary Figure 1).

### The molecular basis for MAT specificity

To understand the mechanisms that dictate the observed differences in catalytic specificity between hMAT2A and eMAT, we investigated three alternative explanations: *(i)* different oligomeric states; *(ii)* residue differences in the substrate binding pockets; *(iii)* different conformational sampling/protein dynamics, caused by sequence differences in regions remote from the active site.

#### *(i)* Oligomeric state

The active sites of both enzymes are located at the dimer interface^31, 32^. Accordingly, we investigated whether differences between the native oligomeric states of either eMAT or hMAT2A underlie their different substrate specificity. We used the NTP analogues adenosine-5’-[(β, γ)-imido]triphosphate (AppNHp), guanosine-5′-[(β, γ)-imido]triphosphate (GppNHp), cytidine-5′-[(β, γ)-methyleno] triphosphate (CppCp), and uridine-5’-[(β, γ)-imido]triphosphate (UppNHp)], since the MATs can catalyze the transferase step of the reaction to yield SNM analogues, but triphosphate hydrolysis cannot proceed and the product-bound state is thus trapped in the catalytic binding pocket (at least for the timescale of these experiments)^33^.

In mammalian cells, hMAT2A exits as heterotetramer constituted by an hMAT2A homodimer, which forms the catalytic unit, and two regulatory subunits hMAT2β^34^. Since the enzyme catalytic pocket is at the hMAT2A homodimer interface and the regulatory subunits are not required for catalysis^35^, we here focus our study on the hMAT2A homodimer. We confirmed using size exclusion chromatography that native apo-hMAT2A exists in equilibrium between monomeric (63%) and dimeric (37%) states (Figure 2a), whereas native apo-eMAT is tetrameric (Figure 2b). The oligomeric equilibrium of hMAT2A shifts almost entirely towards the dimeric state upon incubation with the NTP analogues and methionine (Figure 2a). If any of a nonhydrolyzable NTP, methionine, triphosphate or SAM were added alone (i.e., if the ternary Michaelis complex is unable to form), no change in the oligomeric state was observed (Supplementary Figure 3a). This result suggests that formation of the ternary Michaelis complex (enzyme:NTP:Met) drives dimer formation in the case of hMAT2A. In the case of eMAT, no change in oligomeric state was observed (Figure 2b and Supplementary Figure 3b). Because there were no observed differences between the cognate and non-cognate analogs with either enzyme, it can be concluded that the differences in substrate specificity are independent of the oligomeric state of the enzymes.

**Figure 2.**
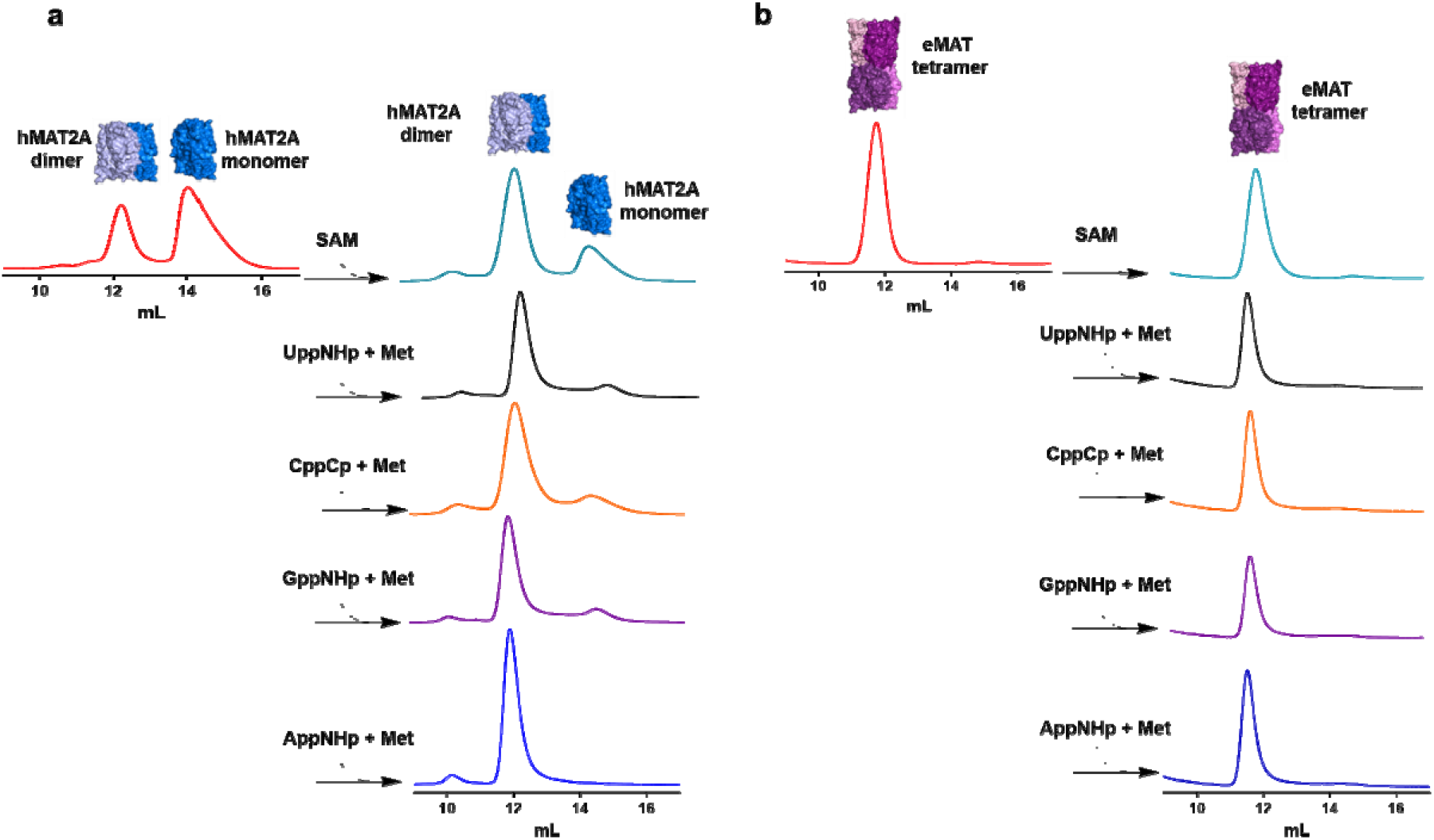
Analysis of oligomeric state of hMAT2A and eMAT by size exclusion chromatography. **a)** hMAT2A (20 μM) is incubated with nonhydrolyzable NTPs (1 mM) adenosine-5’-[(β, γ)-imido]triphosphate (AppNHp), guanosine-5′-[(β, γ)-imido]triphosphate (GppNHp), cytidine-5’-[(β, γ)-methyleno] triphosphate (CppCp), uridine-5′-[(β, γ)-imido]triphosphate (UppNHp)] together with methionine (Met, 10 mM) using reaction buffer (100 mM HEPES, 10 mM KCl, 10 mM MgCl_2_ C for 1 hr). hMAT2A is in an equilibrium of a monomer and dimer. When incubated with both substrates convert completely in dimeric state. No change in oligomeric state when incubated with SAM. **b)** eMAT (20 μM) is incubated using the same condition as used for hMAT2A. eMAT is in a tetrameric state and no change in oligomeric state after incubating with both substrates and SAM was observed. Size exclusion chromatography was performed using GE Healthcare Life Sciences using Superdex 200 Increase 10/300 GL column.

#### *(ii)* The substrate binding site

To investigate the structural basis for substrate promiscuity, we then solved structures of eMAT and hMAT2A in complex with various substrates and substrate analogues (Supplementary Table 1). The structure of hMAT2A in complex with SAM and imidotriphosphate (PPNP) has previously been reported^32^ (Figure 3a). Here we solved a crystal structure of eMAT in the presence of ATP and methionine, which enabled us to capture the SAM product-bound state of eMAT at a resolution of 1.95 Å (Figure 3b). This allowed us to align the eMAT: SAM structure to the previously published hMAT2A: SAM structure (Figure 3c). The active site structures of eMAT and hMAT2A were essentially identical; the only difference was that the eMAT structure has pyrophosphate (PPi) and orthophosphate (Pi) bound in the lower part of the active site, whereas the hMAT2A structure has PPNP bound in the same position (the substrate in the hMAT2A protein crystal being the analogue AppNHp, rather than ATP). Regarding the nucleoside binding region, we observe stabilizing interactions between the enzyme and adenine ring that include a π-stacking interaction (3.5 Å) with Phe230/250 (eMAT/hMAT2A numbering) and hydrogen bonds between the amine group of the adenine ring and the carbonyl oxygen of Arg229/249 and the N1 adenine nitrogen with the side chain of Thr227/Ser247. Closure of the active site loop brings Ile102/117 close to the adenine ring, forming van der Waals contacts.

**Figure 3.**
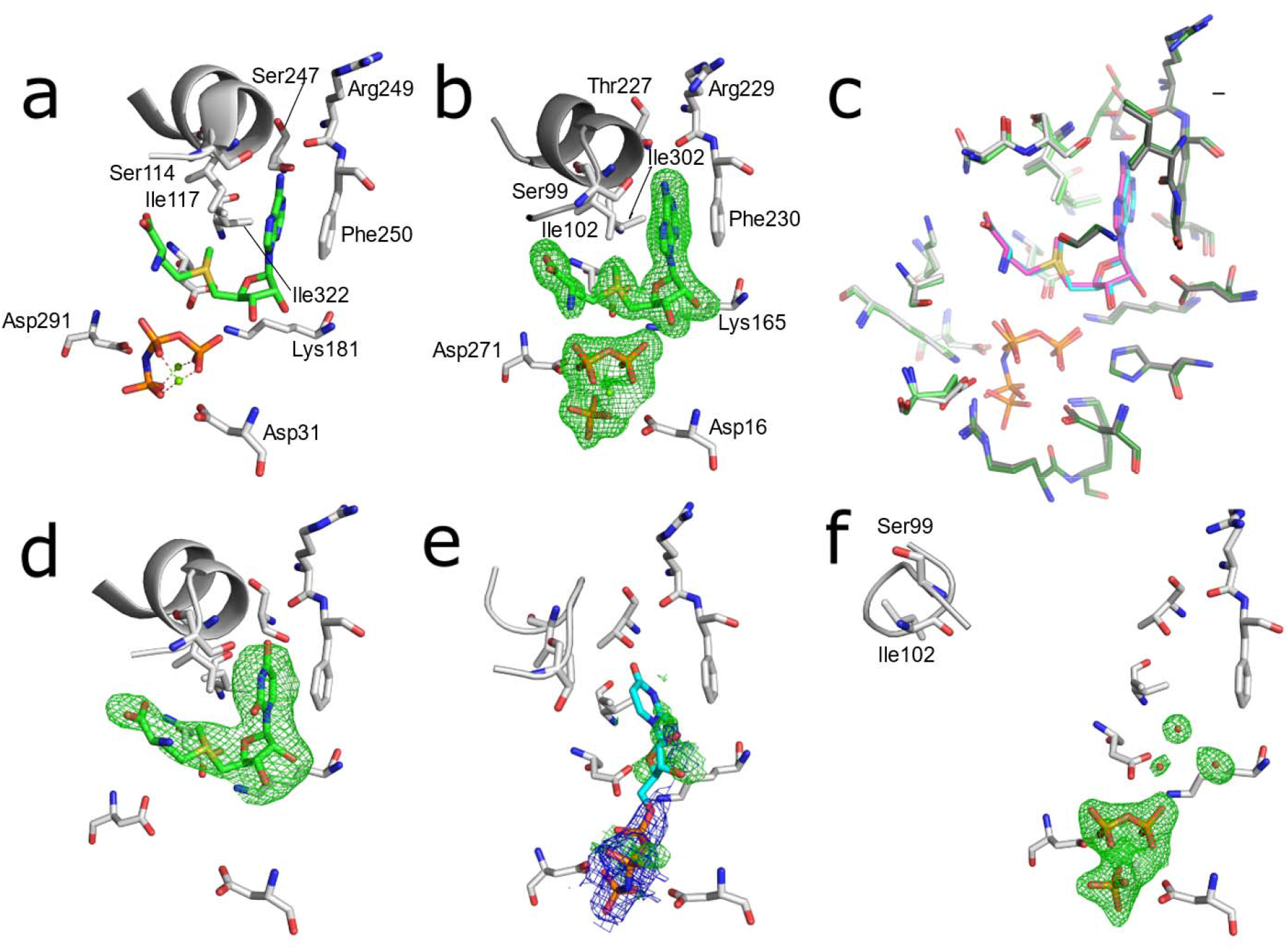
The substrate binding sites of eMAT and hMAT2A with bound substrates. Omit electron density (m*F*_o_-D*F*_c_) is shown in green mesh (3.0 σ), 2m*F*_o_-D*F*_c_ density is shown as blue mesh (1.5σ). (a) The published structure of hMAT2A bound to the products SAM and PPNP (PDB ID 4NDN). (b) Structure of eMAT obtained via cocrystallization with ATP and methionine. (c) A superimposition of the substrate binding sites of eMAT (green, cyan SAM, PPi and Pi) and hMAT2A (grey, magenta SAM, PPNP). The binding site is comprised of two chains within a homodimer; these are distinguished by light/dark colouring. (d) hMAT2A in complex with SUM after cocrystallization with UppNHp. (e) eMAT cocrystallized with UppNHp (2.24 Å); the PPNP group is included in the model and shown with 2m*F*_o_-D*F*_c_, ambiguous omit density potentially corresponding to disordered substrate/product is shown. A poorly fitting model of UppNHp is shown in stick representation (cyan). (f) eMAT bound to the products PPi and Pi (1.89 Å), with the active site loop captured in the “wide-open” conformation obtained *via* cocrystallization with CTP and methionine.

The alignment of the residues within the active site is strikingly similar, both in terms of identity (20/21) and structure (RMSD 0.5 Å). Indeed, every amino acid side chain adopts the same rotamer and the product is bound in an identical conformation. The only difference between the two structures is a conserved substitution at position 227/247 (Thr in eMAT *vs*. Ser in hMAT2A; Supplementary Figure 4). To investigate the effect of this substitution, we made Ser247Thr and Thr227Ser mutants in hMAT2A and eMAT, respectively. Neither mutation resulted in any significant change in substrate specificity in either enzyme (Supplementary Figures 5). Thus, the amino acid composition and structures of the substrate binding sites of the two enzymes do not explain the observed differences in their substrate specificity.

To better understand the structural basis for catalytic promiscuity in hMAT2A, we co-crystallized the enzyme with the non-cognate substrate UTP analog, UppNHP, and solved the structure to 2.5 Å resolution. Imido-NTP analogs have been used as partial inhibitors of MAT because, while they are still substrates for the first methionine transfer step and form the various SNM products, the imido-linkage between the β–γ phosphate units prevents hydrolysis of the triphosphate moiety^33^. Triphosphate hydrolysis is thought to provide energy for active site loop (residues 113-131 in hMAT2A, 98-108 in eMAT) opening and product release^28, 32^. The structure revealed the product SUM bound at high occupancy within the active site (Figure 3d) and was mostly identical to the structure of hMAT2A in complex with SAM. One difference that is not relevant to the nucleoside/SNM binding region is the observation that the PPNP group is completely absent in the structure. The active site loop was fully closed and interacted with SUM in the same manner as with SAM, and the π-stacking interaction with Phe250 was present. The main difference was that the hydrogen bond between the amine group of the adenine and the carbonyl oxygen of Arg249 is not present, although a hydrogen bond between the carbonyl group of the uridine ring and Ser247 is observed. The close similarity between the hMAT2A:SAM and hMAT2A:SUM complexes is consistent with the similar rates of the UTP and ATP turnover observed in the enzyme kinetics (Table 1), suggesting both substrates are stable in catalytically competed configurations.

In contrast to the eMAT:ATP, hMAT2A:ATP and hMAT2A:UppNHp structures, when eMAT was co-crystallized with non-cognate NTPs (CTP/UTP/GTP) we did not observe electron density for any SNM product. Within the active site of the 1.89 Å resolution CTP co-crystal, clear difference density for the PPi and Pi products was observed (Figure 3), although there is unambiguously no electron density for the SCM product. Ordered water molecules were observable, suggesting the SCM had fully diffused from the active site. The active site loop, which was stable and closed in the SAM structure, was instead observed in a “wide-open” conformation, which we believe is the first time this fully open conformation has been fully modelled. For the co-crystals of eMAT with UTP (2.25 Å) and GTP (2.39 Å), which crystallized in a different space group to CTP (Supplementary Table 1), we again observe PPi and Pi in good electron density (Supplementary Figure 6). Like the CTP co-crystal, we do not observe density for the products of the methionine transferase reaction. In these crystals the density in this region appears to correspond to a phosphate molecule, which has presumably re-bound to the protein after hydrolysis. In these UTP/GTP co-crystals, the active site loop is neither closed, as in the eMAT:SAM structure, nor wide-open, as in the eMAT:CTP co-crystal. Instead, it adopts a disordered intermediate conformation. These results suggest that the non-adenine containing SNM products are less stable within the active site of eMAT than SAM, consistent with eMAT being selective for ATP.

Finally, the co-crystals of eMAT with UppNHp (2.24 Å) and GppNHp (2.50 Å) displayed clear electron density for the PPNP group and weaker electron density in the nucleoside binding region. This density is not consistent with re-bound phosphate, as observed in the GTP/UTP co-crystals (Supplementary Figure 6), both because of the non-tetrahedral shape of the density and the observation that there is no way that free phosphate could be present in these crystals, since the substrate analog cannot undergo hydrolysis between the β–γ phosphodiester bond, as for the NTPs. The density also extended continuously from the PPNP moiety (Supplementary Figure 6). Finally, the active site loop was in a partially open, disordered, conformation. The poor electron density for the nucleoside groups in these structures could be due to either (or a combination of) disordered binding of the nucleoside moiety of the substrate or diffusion/disorder of the SUM/SGM product. Even if the weaker density is fully due to substrate turnover and diffusion of SUM/SGM, this behavior is very different to the eMAT:ATP and hMAT2A:UppNHp structures, in which the product was clearly stable within the active site. Thus, in contrast to hMAT2A, which interacts in an essentially identical manner with ATP and the non-cognate substrates, eMAT appears to be selective for adenine-containing substrates, because the adenine containing nucleoside moiety is more stable within the active site, which is again consistent with the enzyme kinetics (Table 1).

#### *(iii)* Differences in protein and substrate dynamics

The crystallographic analysis of the UppNHp/GppNHp:eMAT complexes suggested the poor density could be due, at least in part, to a disordered substrate binding mode. To examine this possibility in more detail, we performed molecular dynamics (MD) simulations of the eMAT tetramer and hMAT2A homodimer, each in complex with both ATP and UTP to investigate whether there were significant differences between enzyme:substrate interactions across proteins that could explain their differing substrate specificities. The methionine substrate was not considered in the simulations. In order not to bias these simulations, all four simulations began with a starting model in which the loop was fully closed over the active site, i.e., the eMAT:UTP bound structure was modelled on the stable SUM bound structure observed in hMAT2A to position the uridine moiety. However, during triplicate 1 μs MD simulations of each complex, (Supplementary Figure 7) the closed conformation was found to be unstable in the absence of bound methionine. Over the course of the simulations, nearly all active site loops across all complexes and replicates transitioned to a dynamic open conformation. To avoid sampling bias arising from the variability in closed-to-open transition time points across domains and replicates, we used representative open-state structures at the endpoint of these trajectories as seed structures for open-state simulations.

Triplicate 500 ns simulations of all four systems (Supplementary Figure 8) show no clear differences in backbone dynamics between eMAT and hMAT2A (Supplementary Figure 9 & 10), suggesting that conformational fluctuations in the protein backbone are not responsible for nucleotide discrimination in eMAT. During these simulations, changes in substrate positioning were also observed (Supplementary Figure 11). While the triphosphate moiety in both ATP and UTP-bound simulations remain stable, the sugar and purine/pyrimidine moieties adopt varied conformations, as the active site loop open state lacks the stabilizing interaction with Ile102’ observed in the closed-state structures (Figures 3; Supplementary Figure 12). The resulting substrate conformations are largely dictated by rotations around the β and χ dihedral angles (Figure 4).

**Figure 4.**
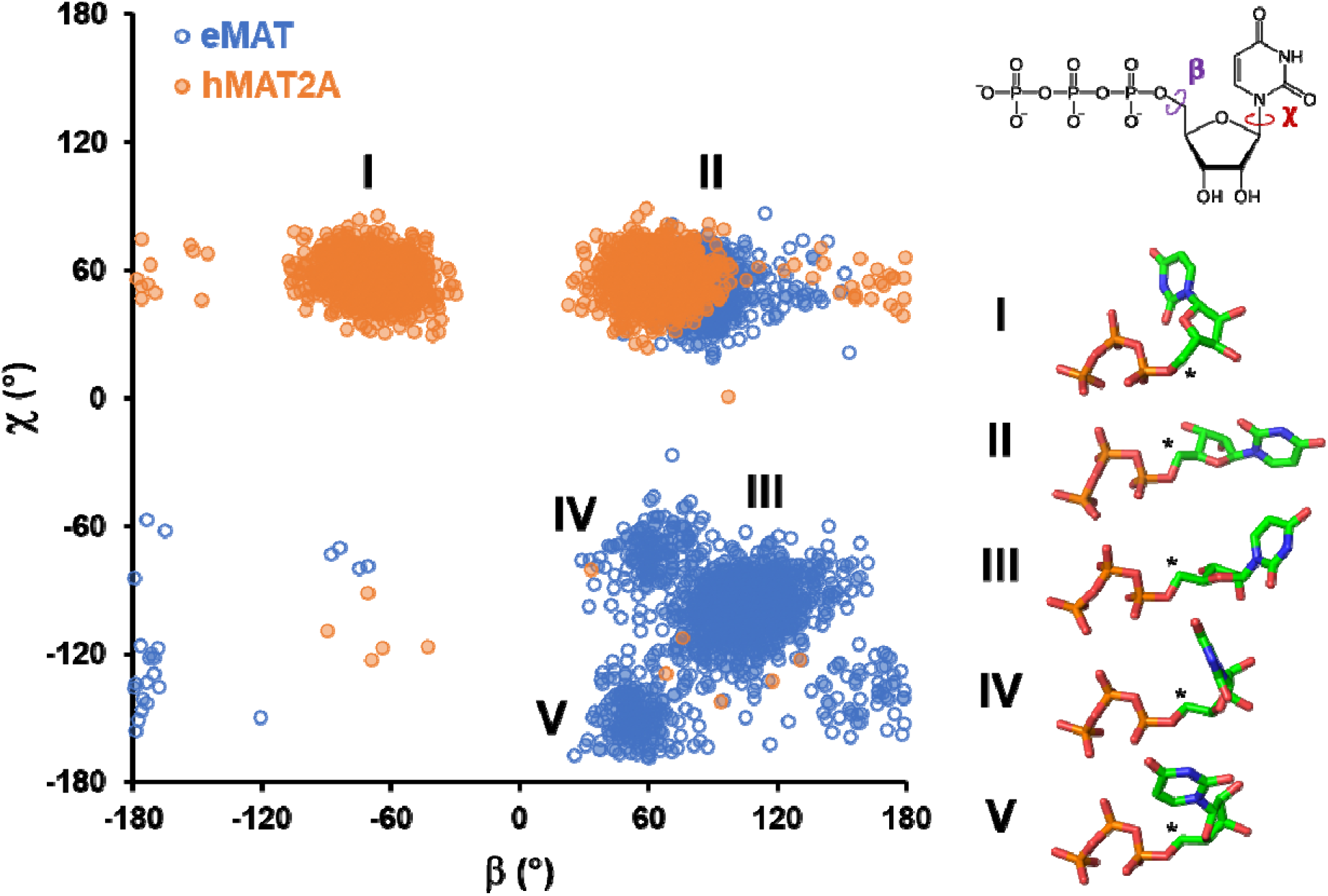
Conformational states populated by the UTP substrate bound to eMAT or hMAT2A. A plot of the UTP β vs χ dihedral angles (shown on the inset 2D representation of UTP) highlights the differences in conformational diversity exhibited by UTP in complex with eMAT or hMAT2A. Each data point represents a dihedral angle pair from one UTP molecule in one simulation frame, sampled every nanosecond over triplicate 500 ns trajectories. Dihedral angle measurements from different enzyme subunits were treated as independent data points. Major conformational clusters in the resulting landscape are shown as a stick representation with the electrophilic C5’ identified with an asterisk. The different conformational states adopted by UTP in eMAT and hMAT2A are indicative of differing enzyme-substrate interactions that constrain the UTP conformation and may contribute to enzyme specificity.

In hMAT2A, the β dihedral angle varies between gauche^□^ (I) and gauche^+^ (II) conformations, while the χ dihedral angle is observed to predominantly adopt the favourable gauche^+^ (II) conformation. In eMAT however, interactions between Lys165 and the O4’ / O5’ of UTP prevent the adoption of the β dihedral gauche^□^ conformation. In several domain replicates of the eMAT:UTP complex, a strained eclipsed χ dihedral conformation (III) is observed that likely arises from electrostatic repulsion between UTP’s uracil moiety and the nearby Asp118 side chain and Gly117 backbone carbonyl (Supplementary Figure 13). Dissociation of the nucleotide in eMAT:UTP complex was never observed over the timescale of the simulations performed due to strongly favourable interactions between bound Mg^2+^ ions and the triphosphate moiety^36, 37^

However, the strained nucleotide conformations observed in the eMAT:UTP complex, which are absent in the hMAT2A:UTP complex, may indicate a weaker binding capacity for UTP in the open state for eMAT comparatively to hMAT2A. Interestingly, in one instance, this conformation also transitioned to an alternate conformation that rapidly fluctuated between gauche^□^ (Figure 4, IV) and trans (Figure 4, V) χ dihedral conformations. The trans (V) χ dihedral conformation, which is observed only in the eMAT:UTP complex, positions the UTP pyrimidine ring such that it blocks binding and nucleophilic attack from methionine on C5’ of UTP (Figure 4). Thus, subtle substrate-enzyme interactions in eMAT that are not present in hMAT2A result in an altered UTP conformational landscape that destabilizes substrate binding and forces the adoption of non-productive binding modes. These observations are consistent with the disorder in the active site loop and the poor electron density for the nucleoside moieties in the substrates/products from co-crystallization with CTP/UTP/GTP/UppNHp/GppNHp (Figure 3), as well as the observed substrate specificity of eMAT (Table 1). It is however important to note that these results show that productive binding of non-cognate NTPs can occur (consistent with slow turnover), but that they are less stable/frequent in eMAT than in hMAT2A.

### The promiscuity of hMAT2A is relevant *in vivo*

Having established that hMAT2A is promiscuous, and eMAT is specific we compared the reported the physiological concentrations of these NTPs in human^38^ and *E. coli*^39^ cells (Supplementary Table 2): in human cells, the concentration of ATP is ~2.5 mM and the other NTPs (GTP 0.2 mM, CTP 0.08 mM and UTP 0.2 mM) are almost 10-fold lower, whereas in *E. coli* the NTPs are all present at similar concentrations.

We then investigated whether the promiscuous products of hMAT2A could be detected *in vivo*. We performed metabolite analysis of SNM abundance using liquid chromatography-mass spectrometry (LC-MS) of extracts from the normal human liver cell line THLE-2 and the hepatocarcinoma cell (HCC) line HepG2, in which hMAT2A is known to be upregulated^40–42^. Notably, we observed the breakdown product of SGM, methionine thioguanosine (MTG), in the HepG2 cell line (Figure 5a) and not in a THLE-2 (Supplementary Figure 14). As a control, we could detect SAM and MTA in both the samples (Appendix). To the best of our knowledge, there is no other way to form MTG other than from SGM (Figure 5b). The presence of MTG was confirmed by mass spectrometry analysis (Figure 5c). It is unclear whether MTG formed during the extraction procedure or is generated endogenously in the cells. However, SNM analogs were found to have comparable stability in aqueous buffer over the same period. (Supplementary Figure 15), suggesting SGM is not significantly less stable and more prone to degradation. Even though the *K*_M_ of hMAT2A with CTP (0.08 mM) is lower than for ATP (0.27 mM) we did not detect SCM or MTC analogs in any of the cell lines within the sensitivity range of the experiment; most likely due to the lower CTP concentration within the cells (0.083 in normal cells and 0.4 mM in cancer cells). It is currently unclear what role the hMAT2β subunit has on specificity, although the work presented here indicates specificity is primarily dictated by the catalytic subunit.

**Figure 5.**
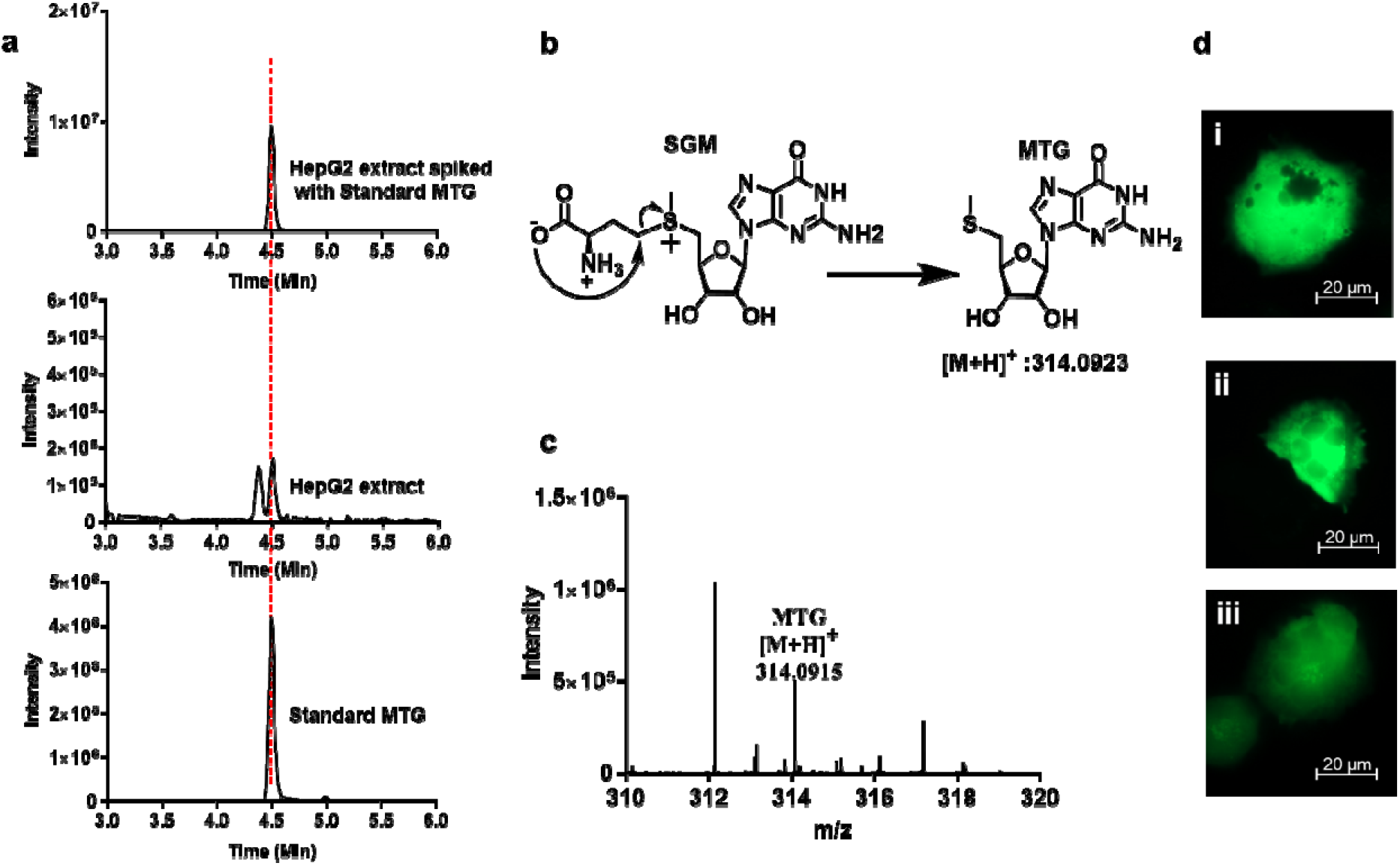
LC-MS analysis of metabolite and effect of SGM, SAM on HepG2. **a)** Extracted chromatograms of the standard MTG, HepG2 cell extract, cell extract samples spiked with the standard MTG. **b)** Schematic representation of degradation of SGM in to MTG after attack of carboxylate on the γ carbon atom of the methionine. **c)** Mass spectrum of HepG2 extract showing the mass of MTG [M+H]^+^ 314.0915. Data was collected using a Q-Exactive HF mass spectrometer coupled with Waters UPLC ACQUITY M-Class liquid chromatography system. An analytical column (ACQUITY UPLC HSS T3 1.8 um, 1.0 × 150 mm) was used for sample chromatographic separation. **d)** Fluorescence microscopy images showing no morphological effect of SGM and SAM on HepG2 cells. HepG2 cells electroporated with (i) pmaxGFP plasmid and with (ii) 1 mM SAM, (iii) 1 mM SGM. Imaging is done using Celldiscover 7 microscope with 20X resolution with 2× magnification changer. Experiment was performed in biological triplicate.

After the inferred identification of SGM in liver cancer cells, we investigated whether SGM resulted in cellular toxicity or any cellular morphology changes. To overcome the low cell membrane permeability of SGM,^43, 44^ we performed cell electroporation in the presence of three different concentration of SGM (0.01 mM, 0.1 mM, and 1 mM). Electroporation was carried out along with a pmaxGFP plasmid to allow fluorescence microscopy observation and possibly detect any morphological changes. Cells were observed after an overnight incubation. The number of cells in the sample electroporated with SGM was comparable to the control (electroporation only with pmaxGFP plasmid) (Supplementary Figure 16a), even at the highest SGM concentration, which indicates that the concentrations of SGM that were used do not affect cell survival. No microscopic effects on cell morphology could be detected for HepG2 (Figure 5d; Supplementary Figure 16b and c) nor any change in the cell survival. The same experiment was performed using different concentration of SAM (0.01 mM, 0.1 mM, and 1 mM), for HepG2 (Figure 5d; Supplementary Figure 16d and e) resulting in the same observations. Since SGM carries the same methyl transferring group as SAM, it is possible that SGM can neutrally substitute for SAM in the methylation or polyamine downstream pathways (Supplementary Figure 17), or that it is simply inert. Overall, we have shown that hMAT2A produces SGM *in vivo*, that SGM (and/or its breakdown product MTG) is present in the cancer cell line HepG2 in which hMAT2A is upregulated, suggesting that it could be potentially used as a biomarker, and that SGM is not toxic for human cells within the parameters of this experiment.

## Discussion

The enzyme kinetics and structural analysis suggests that the catalytic specificity of eMAT is a result of the non-cognate substrates failing to adopt stable and catalytically competent binding modes. This leads to two questions: first, why are the unique substrate binding modes observed in the eMAT:UTP simulations not catalytically competent? The sulfur of methionine performs its nucleophilic substitution at the ribose C5’ atom; thus, the accessibility of this atom is of paramount importance. In the non-productive states sampled by UTP throughout the simulation, the position of the C5’ atom is sterically occluded by the pyrimidine ring and methionine attack is sterically blocked (Figure 4). Clearly, UTP is a viable substrate for eMAT; indeed, we observe in the MD simulations that catalytically productive enzyme:substrate complexes are stable for hundreds of nanoseconds. Thus, the disorder observed here is best conceptualized as a partial depletion of catalytically productive substrate binding and weaker binding stability, compared with the cognate substrate, ATP. Second, what are the contributions of structural dynamics to hMAT2A catalysis with non-cognate substrates in comparison to eMAT? The active sites are essentially identical (20/21 residues) and substitutions of Ser/Thr in either enzyme at the one variant position have no effect on specificity. However, there are many sequence differences between eMAT and hMAT2A in the second and third shells of the active site loop (Supplementary Figure 18). A plausible explanation is therefore that the crystallographic closed state of hMAT2A observed in the presence of non-cognate substrates is promoted by additional stabilizing interactions in the second and third shells of the active site loop, even though non-cognate substrates make fewer stabilizing interactions with first shell residues. In contrast, eMAT cannot as easily sustain the closed active site loop conformation without the additional stabilizing interactions from the adenine group, which are not present in the binding modes of the other non-cognate nucleotides.

The selective pressure that drove the divergence in catalytic specificity between these orthologous enzymes most likely relates to the different cellular abundance of these molecules, i.e., there has been little selective pressure for hMAT2A to be specific because the other NTPs are not present at sufficiently high concentrations to compete with ATP. Indeed, the concentration of ATP is ~10-fold higher than the *K*_M_, whereas for GTP/CTP/UTP the physiological concentrations are at or below the respective *K*_M_ values (Table 1). In contrast, the concentrations of these NTPs in *E. coli* are more similar: ATP is 3.5 mM while GTP is 1.6 mM. Thus, eMAT likely evolved specificity owing to the selective pressure to discriminate between ATP and other nucleotides: the *K*_M_ of eMAT for ATP is at least 16-fold lower than for any of the non-cognate NTPs (Table 1).

Finally, in this work we showed how MAT promiscuity is relevant *in vivo* as putative example of “underground metabolism”^10^. It is thought that promiscuous functions of enzymes are likely to be physiologically irrelevant^1^. For instance, many promiscuous activities cannot occur at sufficiently high frequency to be relevant owing to the substrate concentrations encountered in physiological contexts^45^, or the extremely low catalytic efficiency of many promiscuous activities making it irrelevant on biological timescales^46–48^. This study is therefore a rare example where we could detect the promiscuous activity of hMAT2A for GTP *in vivo*. Moreover, we showed that it could be used as a biomarker to distinguish between normal and cancer cell lines.

In summary, these results show how enzyme dynamics have substantial effects on the conformational sampling of substrates within the active site of an enzyme, which can in turn result in large changes in catalytic specificity. The concept of non-productive substrate binding is not new^49^, nor is the notion that protein dynamics can affect substrate turnover^50, 51^, but this is an interesting example where the link between these two effects can be clearly seen. Moreover, because we have compared orthologous enzymes that have been on different evolutionary trajectories because of their distinct cellular environments, we have been able to show that the sequence differences controlling this specificity originate in the outer shells of the active site, which builds on a growing body of work that supports a model in which these outer-shell residues are critical for maintaining the optimum active site architecture and controlling conformational changes that are important in the catalytic cycle^15^. Consideration of these effects should aid enzyme engineers, evolutionists and synthetic chemists in the design and study of enzymes, substrates, and inhibitors. For example, we hope that this work will aid in the design of SAM analogues with unnatural bases; such analogues could show promise for reaching cellular bio-orthogonal probes or inhibitors of methyltransferases.

## Methods

### Protein expression and purification

The eMAT plasmid was a generous gift from Prof. Ronald E. Viola. *E. coli.* BL21 (DE3) cells were transformed with the eMAT plasmid and protein was expressed as reported previously^19^. Cell pellets were resuspended in lysis Buffer (40 mM Tris-HCl pH 8.0, 300 mM NaCl, 10 mM imidazole) supplemented with 0.5 units turbonuclease (T4330, Sigma Aldrich), 0.3 mg.ml^−1^ lysozyme, 0.2 mM PMSF and 5 mM DTT. Solubilised pellets were lysed by sonication and centrifuged at 30, 000 × g for 30 min. The soluble fraction was applied to a 5 mL HisTrap HP Ni^2+^-NTA IMAC column (GE Healthcare) pre-equilibrated with lysis buffer and washed with 50 mM imidazole. eMAT was eluted in lysis buffer supplemented with 400 mM imidazole and concentrated with an Amicon Ultra-15 spin concentrator (30 kDa MW cut-off, Millipore). eMAT was further purified by size exclusion chromatography (SEC) using a HiLoad® 26/600 Superdex 200 pg column (GE Healthcare) in SEC Buffer A (50 mM Tris-HCl pH 8.0, 100 mM NaCl and 5 mM DTT). Analysis of MAT protein purity was verified with Coomassie SDS polyacrylamide gel electrophoresis and protein concentrations were calculated using the molar extinction coefficient predicted by the ExPASY ProtParam server tool at A_280_. The hMAT2A plasmid was gift from the Jon S. Thorson and purified as reported^52^. hMAT2A pellets were processed in the same manner as eMAT, using sonication and Ni^2+-^ NTA IMAC except for the composition of lysis buffer (50 mM Na_2_HPO_4_ pH 8.0, 300 mM NaCl and 10 mM imidazole). hMAT2A elution was then incubated with 10 mM L-methionine, 10 mM MgCl_2_ and 100 μM UppNHp for 1h on ice before purification in SEC Buffer B (25 mM HEPES pH 7.6, 150 mM NaCl, 5 mM KCl, 5 mM DTT and 10% (v/v) glycerol) for crystallization.

### Protein crystallization, data collection and structure determination

eMAT crystals were grown at 19 °C using the hanging-drop vapour diffusion method with reservoir solutions containing 0.1 M BIS-TRIS pH 6.5 and 10-20% (v/v) ethylene glycol while screening two different lengths of polyethylene glycol (PEG) at varying concentrations: PEG 8000 from 6-9 % (w/v) and PEG 3350 from 16-22 % (w/v). Drops were setup at 1:1 ratio and 1:2 ratio of reservoir to protein volume. Co-crystals formed within 2-4 days at 19 °C with various substrates. hMAT2A-UppNHp was concentrated to 10 mg.ml^−1^ for protein X-ray crystallographic studies. hMAT2A-UppNHp hanging drops were grown at 19 °C at a 1:1 and 1:2 ratio of reservoir to protein volume. The optimised screening matrix consist of 0.1 M BIS-TRIS pH 6.5 and 10% (v/v) ethylene glycol while screening PEG 3350 at concentrations of 7-10% (w/v). Cubic diamond crystals formed within 2 days at 19 °C. The co-crystals were cryoprotected in solutions containing the mother liquor and increasing the concentrations of PEG 8000 or PEG 3350 to 25-35% (w/v) before being flash-frozen in liquid nitrogen. Diffraction data was collected on the macromolecular crystallography beamline (MX2) at the Australian Synchrotron using the Eiger X 6M detector at a wavelength of 0.9537 Å^53^. Data was processed using XDS^54^ and Aimless^55^, and molecular replacement was performed using Phaser^56^. Iterative cycles of manual model building and refinement were performed using Coot 0.9.3^57^ and phenix.refine^58^. Iterative cycles of manual model building and refinement were performed using Coot 0.9.3, and phenix.refine^58^. TLS refinement was used in all cases, using TLS groups automatically selected by phenix.refine. Notably, chain B in the hexagonal space groups exhibited significant disorder in places. All crystallization conditions, data collection and refinement details are provided in Supplementary Table 1.

### Molecular dynamics simulations

All molecular dynamics simulations were carried out using GROMACS 2018.3^59^. Closed-state simulations were run using nucleotide-bound models derived from hMAT2A: SAM, hMAT2A: SUM, and eMAT: SAM crystal structure as starting points. For eMAT:UTP simulations, UTP was modelled in the eMAT:ATP model structure at the ATP position. Open-state simulations were run using the final frame from a randomly selected closed-state simulation replicate in which all domains had transitioned to the open state as starting points. Completed structures were solvated in a dodecahedral simulation box with a minimum distance of 10 A□ from any protein atom to the box wall, followed by addition of roughly 50 mM NaCl into the aqueous phase, neutralizing the system charge. All systems were subjected to steepest-descent energy minimisation followed by a 100 ps equilibration in the NVT ensemble with position restrains of 1000 kJ/mol/nm2 on all protein atoms, with velocities initialising from a Maxwell distribution at 300 K. All NVT equilibrated systems were then subjected to 100 ps equilibration in the NPT ensemble with position restraints of 1000 kJ/mol/nm2 on all protein atoms. Position restraints were released, and free simulation performed at 300 K for 1 μs for each replicate. All simulations were performed using the CHARMM36-feb2021 forcefield^60^. Water was explicitly modelled using the TIP3P model. Ionisable residues were set to their standard protonation state at pH 7. All equilibration and production simulations were conducted under periodic boundary conditions. Temperature was maintained close to the reference value of 300 K using V-rescale temperature coupling. Pressure was maintained close to the reference value of 1 atm using a Parinello-Rahman barostat with isotropic pressure coupling. The LINCS algorithm^61^ was used to constrain the lengths of all bonds to hydrogen. The Verlet cut-off scheme was used to evaluate the non-bonded interaction pair lists. Van der Waals interactions were evaluated using a simple cut off scheme with a radius of 12 A□. Coulomb interactions were evaluated using the Particle Mesh Ewald (PME) method with a grid spacing of 1.6 A□. A 2-fs time-step was used for integrating the equations of motion. GROMACS tools^59^ were used for correction of periodic boundary conditions. Visual Molecular Dynamics (VMD)^62^ was used to view trajectories and for RMSD, RMSF, and dihedral angle calculations, and PyMOL (The PyMOL Molecular Graphics System, Version 2.0 Schrödinger, LLC.) was used to produce figures.

### Mutagenesis

Site directed mutagenesis for Ser247Thr mutation on hMAT2A plasmid and Thr227Ser mutation on eMAT plasmid was carried out using Q5 Site-Directed Mutagenesis Kit (NEB) by following kit protocol and expressed, purified as hMAT2A and eMAT, respectively. The primers used for mutagenesis listed in Supplementary table 3.

### Kinetics Assay for MATs

To observe the reaction efficiency of SNM product formation during catalysis with different substrates ATP/GTP/CTP/UTP (5 mM) and methionine (10 mM), HEPES (100 mM), MgCl_2_ (10 mM), KCl (50 mM) and hMAT2A/eMAT/Ser247Thr hMAT2A/Thr227Ser eMAT (20 μM) were mixed in water, pH was adjusted to 8 with 10% NaOH. The reactions were incubated at 37 □C for 1 hr. Reaction was quench by acetonitrile followed by centrifugation at 12,000 RPM for 5 min to precipitate the enzymes. Finally, supernatant was filtered through 0.22 μm filter (Merck) and injected in UPLC for analysis (Waters UPLC Acquity H class). Diluted reaction aliquots were analyzed by using a HILIC column (SeQuant ZIC-cHILIC 3 μm,100 Å 150 × 2.1 mm PEEK coated HPLC column). An isocratic method was used with solvent A (100 mM ammonium acetate, pH 5.3) 35% and solvent B (acetonitrile) 65% for 15 min. Each injection was 3 μL with a flow rate of 0.3 mL/min and detected at 260 nm. Using this UPLC method retention times for molecules were MTA 1.3 min, MTU 1.3 min, MTC 1.4 min, MTG 1.5 min, adenine 1.6 min, uracil 1.6 min, cytosine 1.8 min, guanine 2 min, SAM 4.1 min, SCM 4.6 min, SUM 4.6 min, SGM 5.3 min, ADP 5.3 min, UDP 6 min, CDP 6.1 min, GDP 6.3 min, ATP 7.5 min, GTP 7.8 min, CTP 8.3 min. Product formation was further confirmed by mass analysis (Appendix). SNM were purified using above mentioned UPLC method and standard curves were plotted. For kinetic assay concentrations of the NTPs were in the range of 0.0251-5 mM and constant methionine concentration 10 mM were used. The kinetic parameters were determined using the Michaels-Menten equation using GraphPad Prism 7.02. The release of nucleotide bases from SNM analogs were also detected by UPLC (Supplementary Figure 5a). SAM is prone to alkaline depurination^63^ but release of nucleotide bases for pyrimidine ring in our reaction conditions might be due to deprotonation at C-5′ in basic conditions followed by the opening of the ribose ring which eliminates nucleotide base, further attack of water reforms ribose ring to give S-ribosylmethionine^64^. Elimination of nucleotide bases was not observed from NTPs (Supplementary Figure 2b) under the same conditions, which demonstrate that release of nucleotide base was from SNM analogs.

### Analytical Size Exclusion chromatography

Size exclusion chromatography was performed using GE Healthcare Life Sciences using Superdex 200 Increase 10/300 GL column. Injection volume was 100 μL, detection at 280 nm and flow rate was 0.5 mL/min. Nonhydrolyzable NTPs (1 mM) Adenosine-5′-[(β,γ)-imido]triphosphate (AppNHp), Guanosine-5′-[(β, γ)-imido]triphosphate (GppNHp), Cytidine-5′-[(β, γ)-methyleno] triphosphate (CppCp), Uridine-5’-[(β, γ)-imido]triphosphate (UppNHp)] were incubated with methionine (L-Met) (10 mM) in HEPES (100 mM), KCl (50 mM), MgCl_2_ (10 mM), pH 8 at 37 □C for 1 hr and then injected in the column.

### Cell culture and extraction of metabolites

HepG2 was grown in DMEM medium containing 10% FBS and penicillin (100 U/ml), streptomycin (100 mg/ml) by incubation in a 5% CO_2_ at 37 □ C with 95% humidity. For routine maintenance, cells were trypsinized and split before becoming fully confluent. Cultured cells were washed with cold PBS (5 mL) twice. Cells (20 M) were harvested by trypsinization using TrypLE Express Enzyme (1X), no phenol red for 3 min at 37 □ C in CO_2_ incubator. Centrifuged for 5 min at 100g. TrypLE was discarded, and pellet was resuspended into cold PBS. Cell pellet was washed with cold PBS twice. Further extraction steps were performed on ice. Internal standards (10 nmol of HEPES and PIPES) were added to sample. Cells were disrupted using 1 mL of cold acetonitrile, methanol, water (40:40:20) with 0.1 M formic acid and glass beads acid washed, by vertexing. Metabolites were collected by the centrifugation. Samples were concentrated using speed vac and finally dissolved in 100 μL of 10% acetonitrile with 0.1% formic acid and filtered through a 0.22 μm filter and injected into LC-MS.

### LC-MS method for metabolite analysis

Data were collected using Q-Exactive HF mass spectrometer (Thermo Fisher Scientific) coupled with Waters UPLC ACQUITY M-Class liquid chromatography system. An analytical column (ACQUITY UPLC HSS T3 1.8 um, 1.0 × 150 mm) was used for sample chromatographic separation. An injection volume of 2 μL was separated at flow rate of 50□μL/min using a gradient of 10–95% solvent B over 8□min, using water with 0.1% formic acid as solvent A and acetonitrile with 0.1% formic acid as solvent B. MS data were collected using Q-Exactive HF mass spectrometer (Thermo Fisher Scientific). The parameters are listed here: spray voltage, 3.0 kV; sheath gas, 16; auxiliary gas, 2; capillary temperature, 250 °C; aux gas heater temp, 150 °C; S-lens RF, 50; tuning method name, HESI; Spray interface, HESI, with metal needle for small flow (1 to 10 μL/min). The mass spectrometry method was set to acquire MS1 data for 14 min, positive mode, mass range 80 to 1,000 m/z. Resolution was set at 60,000. Maximum injection time was 30 ms. Auto gain was targeted to 500000 ions. Extracted ion chromatograms were done using a 5-ppm tolerance and smoothing with Boxcar method using 7 points.

### Cell electroporation with SGM, SAM and pmaxGFP Plasmid

Cells were harvested by trypsinization and 2×10^6^ cells were pelleted by centrifugation at 100g for 3 min. Cells were resuspended in Nucleofector solution from Lonza. SF cell line 4d-Nucleofector X kit S (V4XC-2032) for HepG2 cells. Cells were electroporated with 0.4 μg pmaxGFP plasmid and different concentrations of SGM and SAM (0.01, 0.1, 1 mM) using 4D-Nucleofector X Unit from Lonza. EH-100 program was used for HepG2 by following the manufactures protocol. Cells were incubated overnight in the incubator and observed under the fluorescence microscope. Cells were observed using Leica DMiL microscope using 10X objective. For higher magnification cells were observed using ZEISS Celldiscoverer 7 using 20X objective with 2× magnification changer.

## Author contributions

M.G. performed MAT kinetics assay (including protein purification), ran SEC analysis, performed all the cell experiments (including culturing), metabolites extractions, imaging, processed and analyzed data. M.G., A.V.B ran LC-MS for metabolites, processed and analyzed data. L.L.T. expressed and purified protein for crystallography and collected crystallographic data. L.L.T., A.D., J.S.B. C.J.J. processed, solved, and analyzed crystallographic data. A.D. and M.S. performed molecular dynamics simulations. P.L. and C. J. J. analyzed data and wrote the paper with input from M.G and A. D. P. L., and C.J.J. conceived and supervised the project.

## Acknowledgments

We thank Yohsuke Moriyama for assistance with the cell electroporation experiment and Keiko Kono to share the fluorescent microscope in her Unit. We thank Saacnicteh Toledo-Patino and Benjamin Clifton for insightful comments on this manuscript.

