## supplementary information for "Substrate dynamics contribute to enzymatic specificity in human and bacterial methionine adenosyltransferases"

Madhuri Gade^1^, Li Lynn Tan^2^, Adam M. Damry^2^, Mahakaran Sandhu^2^, Jospeh S. Brock^3^, Andie Delaney^2^, [Alejandro Villar-Briones](https://www.nature.com/articles/s41467-018-05884-0#auth-12)^1^, Colin J. Jackson*^2,4,5^, Paola Laurino*^1^

^1^Protein Engineering and Evolution Unit, Okinawa Institute of Science and Technology Graduate University, 1919-1 Tancha, Onna, Okinawa, Japan 904-0495

^2^ Research School of Chemistry, Australian National University, Canberra, 2601, Australia

^3^ Research School of Biology, Australian National University, Canberra, 2601, Australia

^4^ Australian Research Council Centre of Excellence for Innovations in Peptide and Protein Science, Research School of Chemistry, Australian National University, Canberra, 2601, ACT, Australia

^5^ Australian Research Council Centre of Excellence in Synthetic Biology, Research School of Chemistry, Australian National University, Canberra, 2601, ACT, Australia

**Materials**

ATP, GTP, CTP, UTP, methionine, S-adenosylmethionine (SAM), HEPES, MgCl_2_, KCl, isopropyl-1-thio-β-D-galactopyranoside (IPTG), Tris HCl, Na_2_HPO_4,_ NaH_2_PO_4_, potassium phosphate, NaCl, imidazole, β-mercaptoethanol, dithiothreitol (DTT), kanamycin, glycerol, NaOH, HCl, ammonium acetate, bacto agar, bacto tryptone, bacto yeast extract all other chemicals and HPLC grade solvents were purchased from commercial sources and used as supplied unless otherwise mentioned. PageRuler prestained protein ladder, 10 to 180 kDa and TrypLE Express Enzyme (1X), no phenol red, DMEM - Dulbecco's Modified Eagle Medium, Trypsin–EDTA, Fetal bovine serum (FBS) and PBS, penicillin-streptomycin solution were purchased from ThermoFischer scientific. BL21 (DE3) competent cells and Q5 Site-Directed Mutagenesis Kit were purchased from New England Biolabs (NEB). RedTaq Ready Mix PCR reaction mix and benzonase, complete His-Tag Purification Resin (NiNTA), glass beads acid washed, Bovine Collagen Solution Type I were purchased from Sigma Aldrich. Protein inhibitor cocktail (PIC), bovine serum albumin (BSA) and Lysozyme were purchased from Nacalai Tesque, INC. 12% Mini-PROTEAN TGX Precast Protein Gels, 12-well from BIO-RAD. Amicon centrifugal filters were purchased from Merck. Standards for size exclusion chromatography were purchased from GE Healthcare. AppNHp, GppNHp, UppNHp and CppCp were purchased from Jena Bioscience. THLE-2 cells were purchased from ATCC. BEGM Bronchial Epithelial Cell Growth Medium Bullet Kit was purchased from Lonza. Fibronectin Human Protein were purchased from Life technologies. [Phosphorylethanolamine](javascript:void(0);) was purchased from funakoshi. SF Cell Line 4D-Nucleofector™ X Kit and P1 Primary Cell 4D-Nucleofector™ X Kit S were purchased from Lonza. All the experiments were performed using [**ultrapure water purification system**](https://red.oist.jp/equipment.php?assetkey=RED-00000846) from a MilliQ Integral MT10 type 1 (Millipore).

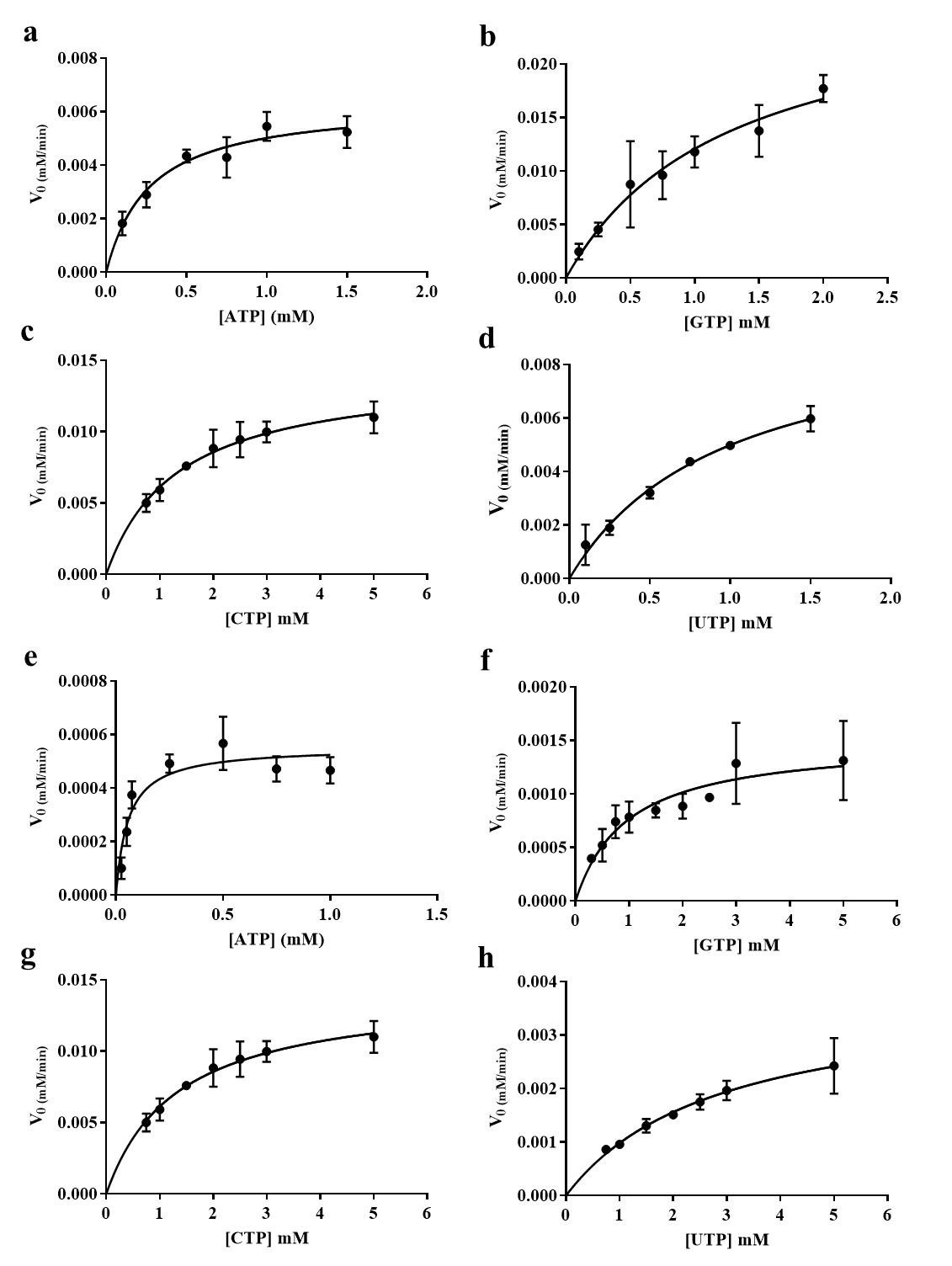

**Supplementary Figure 1. Kinetic characterization of hMAT2A and eMAT using NTPs.** Michael-Mentis plots for hMAT2A forming SAM (a), SGM (b), SCM (c) and SUM (d). Conditions: [hMAT2A] = 0.5 µM; [NTPs] = 0.05 - 2 mM; [methionine] = 10 mM; [HEPES] =100 mM (pH 8, 37 ̊C); [KCl] = 50 mM; [MgCl_2_] = 10 mM. Michael-Mentis plots for eMAT forming SAM (e), SGM (f), SCM (g) and SUM (h). Conditions: [eMAT] = 0.5 µM for ATP, 5 µM for GTP, CTP and 10 µM for UTP. [NTPs] = 0.025 -5 mM. Other reaction conditions are same as hMAT2A. SNM production was analyzed by UPLC, and data fitted to the Michaelis-Menten equation using GraphPad Prism 7.02. Experiments were performed in duplicates and error bars show standard deviation.

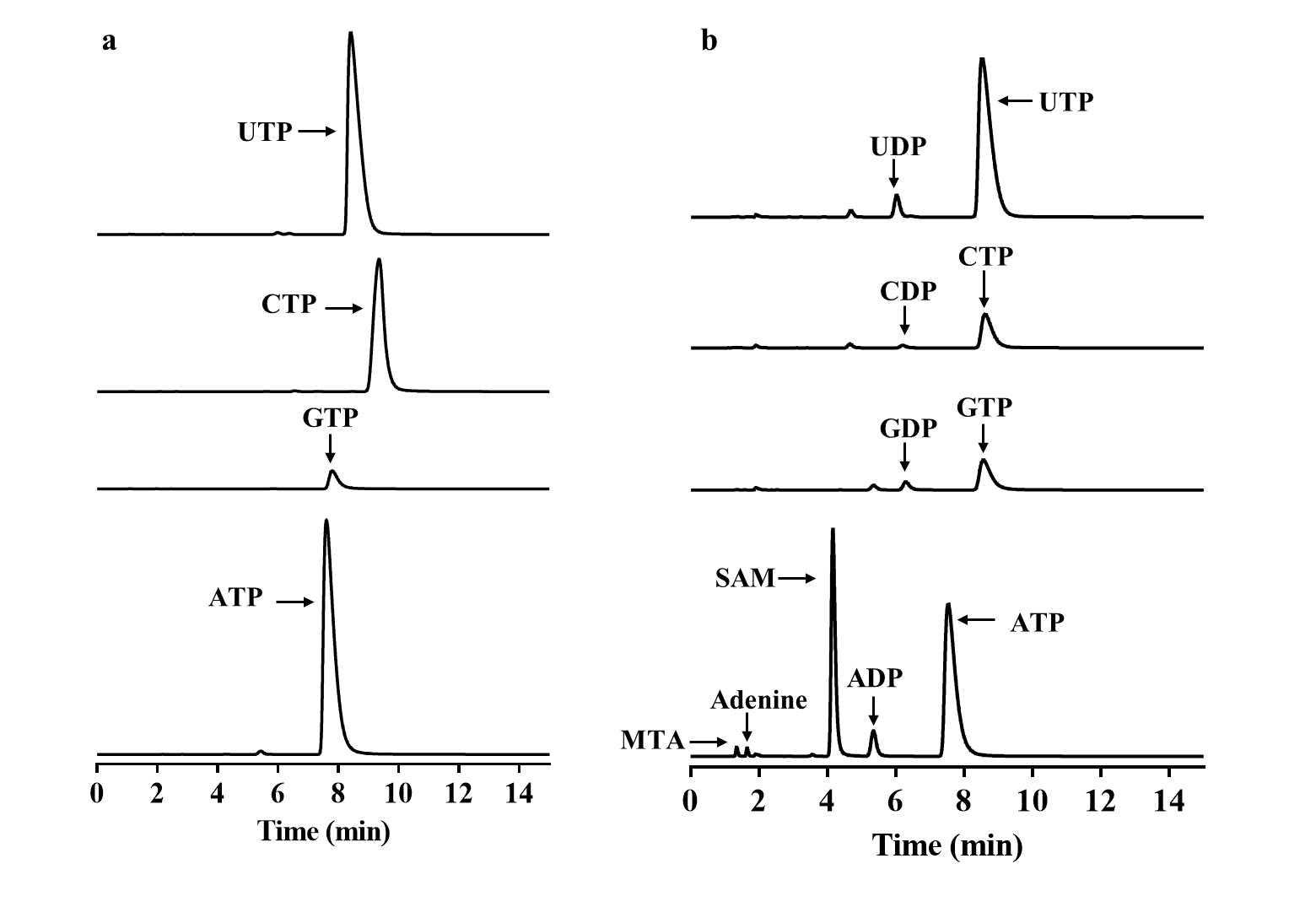

**Supplementary Figure 2. UPLC chromatogram of the reaction between NTP, methionine, without MAT (a) and with eMAT (b).** Reaction details and UPLC method as reported in methods.

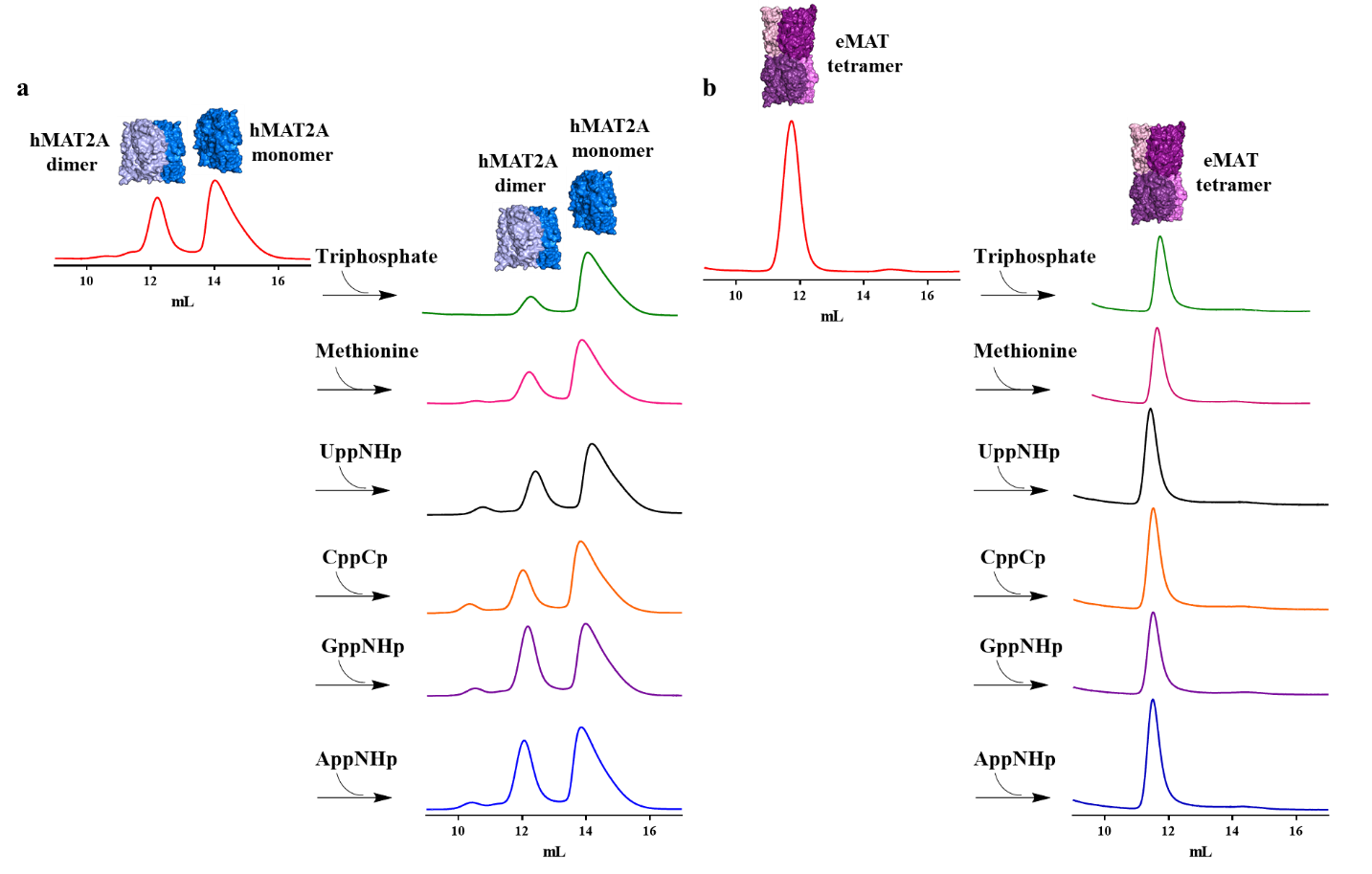

**Supplementary Figure 3. Analysis of oligomeric state of hMAT2A and eMAT by size exclusion chromatography.** hMAT2A is in an equilibrium of a monomer and dimer (a) and eMAT is in a tetrameric state (b). When incubated with nonhydrolyzable NTPs, methionine (Met), triphosphate no change in oligomeric state was observed.

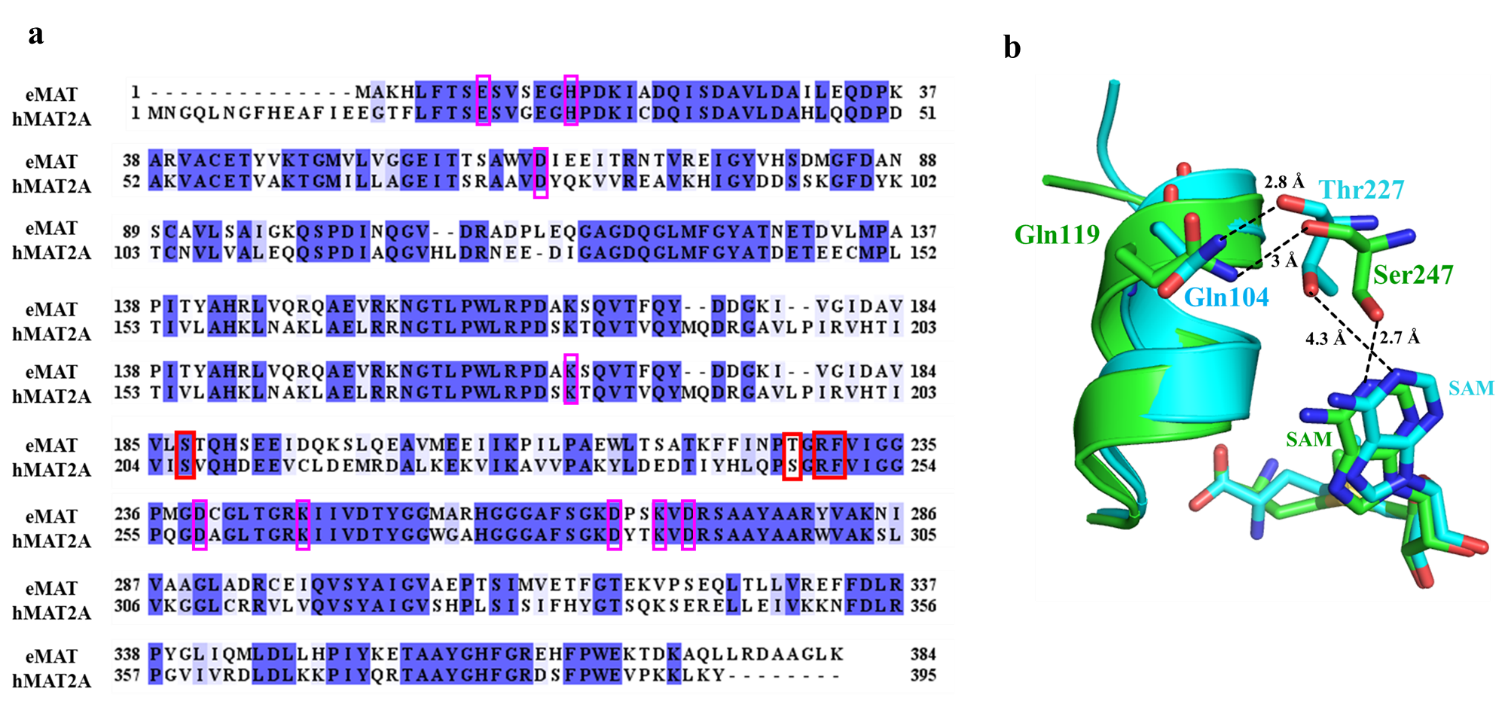

**Supplementary Figure 4.** **Multiple sequence alignment of eMAT and hMAT2A**; **Interaction of gating loop residue Gln with Ser and Thr. a)** Identical amino acid residues are shown in shades of blue. Residues interacting with adenine of SAM are marked in red rectangle. Residues involved in catalysis^5^ are marked in pink rectangle. b) SAM, interacting residue and gating loop is in green color for hMAT2A (PDB ID 4NDN). SAM, interacting residue and gating loop is in cyan color for eMAT (PDB ID 1RG9). N1 of adenine is forming hydrogen bond with Thr227 of eMAT (bonding distance 4.3 Å). N1 of adenine is forming hydrogen bond with Ser247 of hMAT2A (bonding distance 2.7 Å). Backbone of Thr227 and Ser247 is forming hydrogen bond with Gln104 (bonding distance 2.8 Å) and with Gln119 (bonding distance 3 Å) respectively.

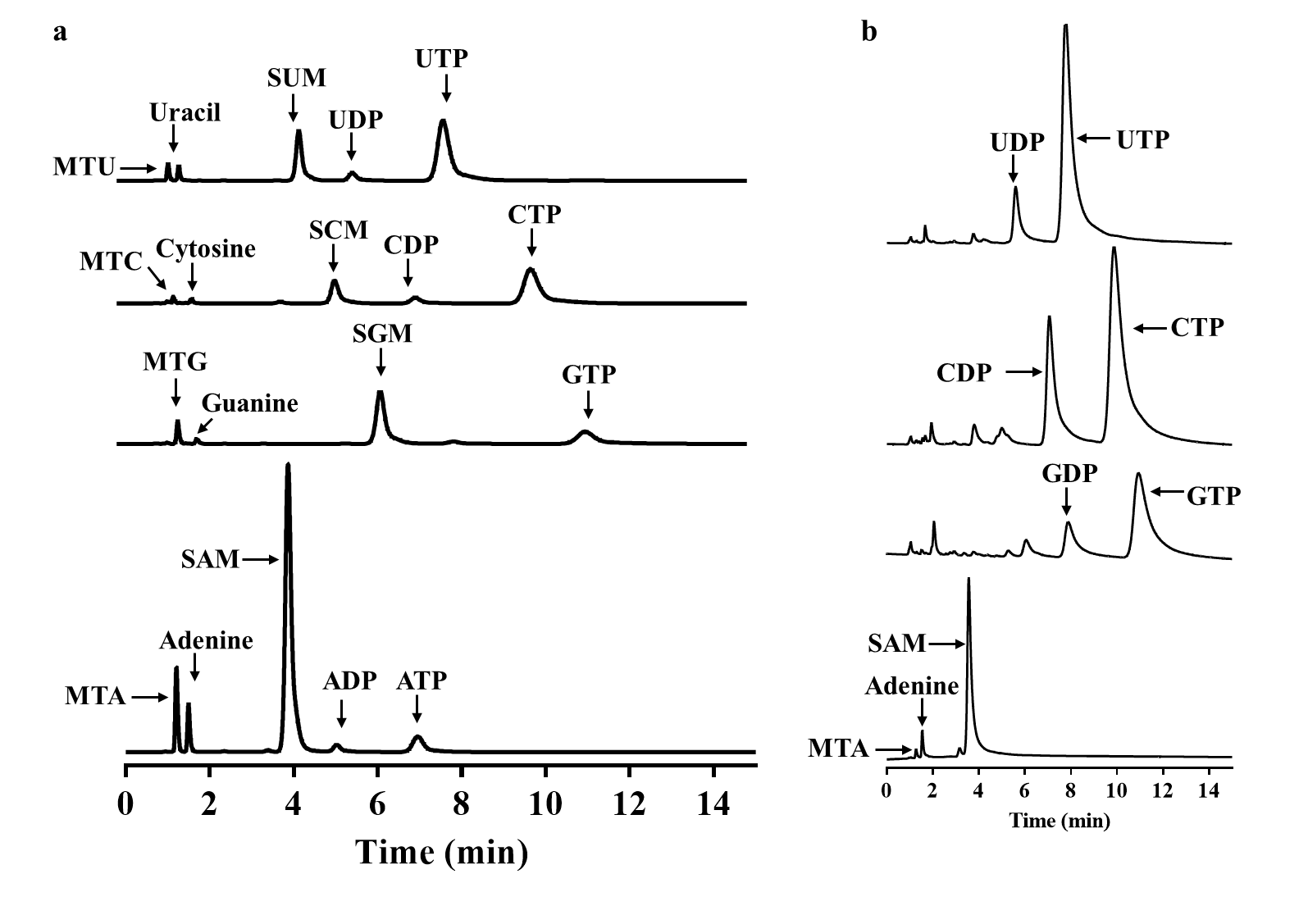

**Supplementary Figure 5. UPLC chromatogram of the reaction between NTP, methionine, and Ser247Thr hMAT2A mutant (a), Thr227Ser eMAT mutant (b).** Reaction details and UPLC method as reported in methods.

**Supplementary Figure 6. Models and electron density for various substrate/product complexes with eMAT.** Omit electron density (m*F*_o_-D*F*_c_) is shown in green mesh (3.0 s), 2m*F*_o_-D*F*_c_ density is shown as blue mesh (1.5 s). (a) eMAT cocrystallized with UTP (2.25 Å); the pyrophosphate and phosphate groups are included in the model and shown with 2m*F*_o_-D*F*_c_, omit density. (b) eMAT cocrystallized with GTP (2.39 Å); the pyrophosphate and phosphate groups are included in the model and shown with 2m*F*_o_-D*F*_c_, omit density. (c) eMAT cocrystallized with GppNHp (2.5 Å); the PPNP group is included in the model and shown with 2m*F*_o_-D*F*_c_, ambiguous omit density potentially corresponding to disordered substrate/product is shown. A poorly fitting model of GppNHp is shown in line representation (magenta). (d & e) Enlarged view of electron density in binding site of UppNHp from two angles showing the density is continuous with the electron density of the PPNP group. (f & g) Omit electron density corresponding to the bound Pi molecule in the eMAT:UTP complex. The density is tetrahedral, and the Pi group is in hydrogen bonding distance to two water molecules and Asp163. This is not unusual as Asp/Glu residues often have elevated pKa values in enzyme active sites.

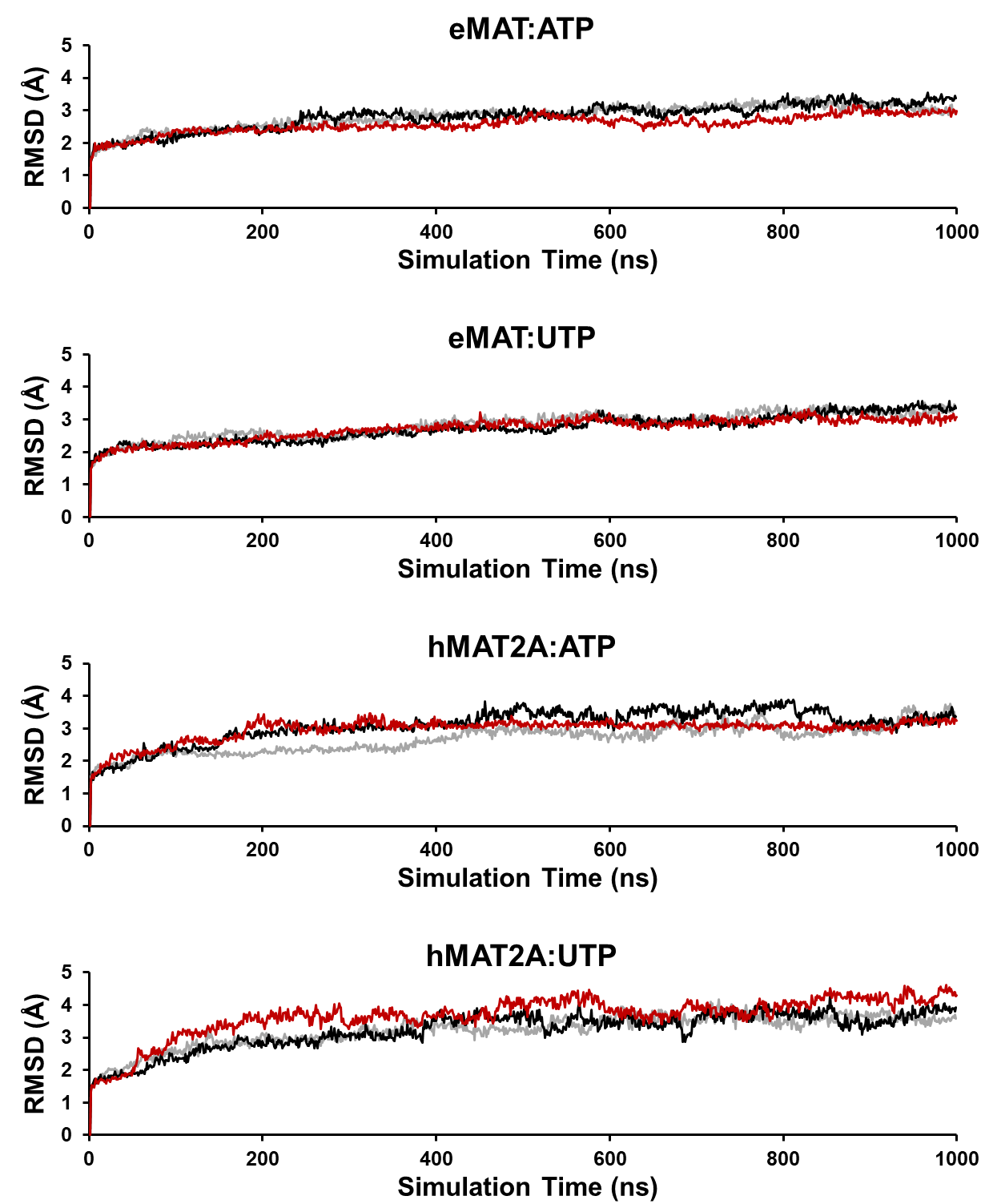

**Supplementary Figure 7. Closed state MD simulation RMSD.** A plot of simulation RMSD versus time for shows a rapid divergence from the closed state input structure followed by slow equilibration that gradually reaches an RMSD plateau, indicative of the slow active site loop opening observed. Line colors indicate simulation replicates for each system modeled and the red line represents the trajectory used as input for open state simulations (Supplementary Figure 8).

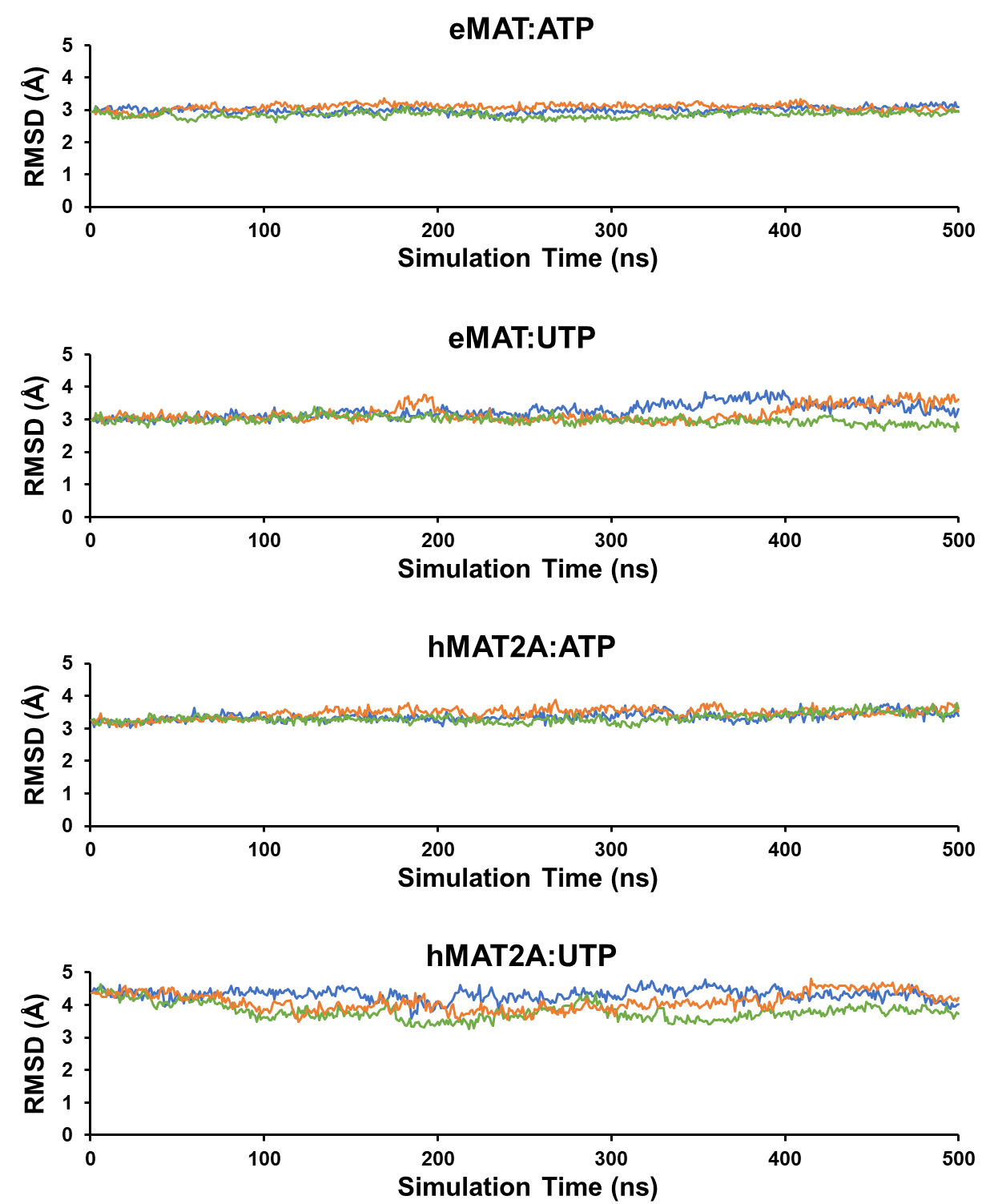

**Supplementary Figure 8. Open state MD simulation RMSD.** A plot of simulation RMSD versus time indicates that open state input structures are already equilibrated by the end of the closed state simulations and remain at a relatively constant RMSD over 500 ns of additional simulation time. The RMSD shown is relative to the closed state input structure. Line colors indicate simulation replicates for each system modeled.

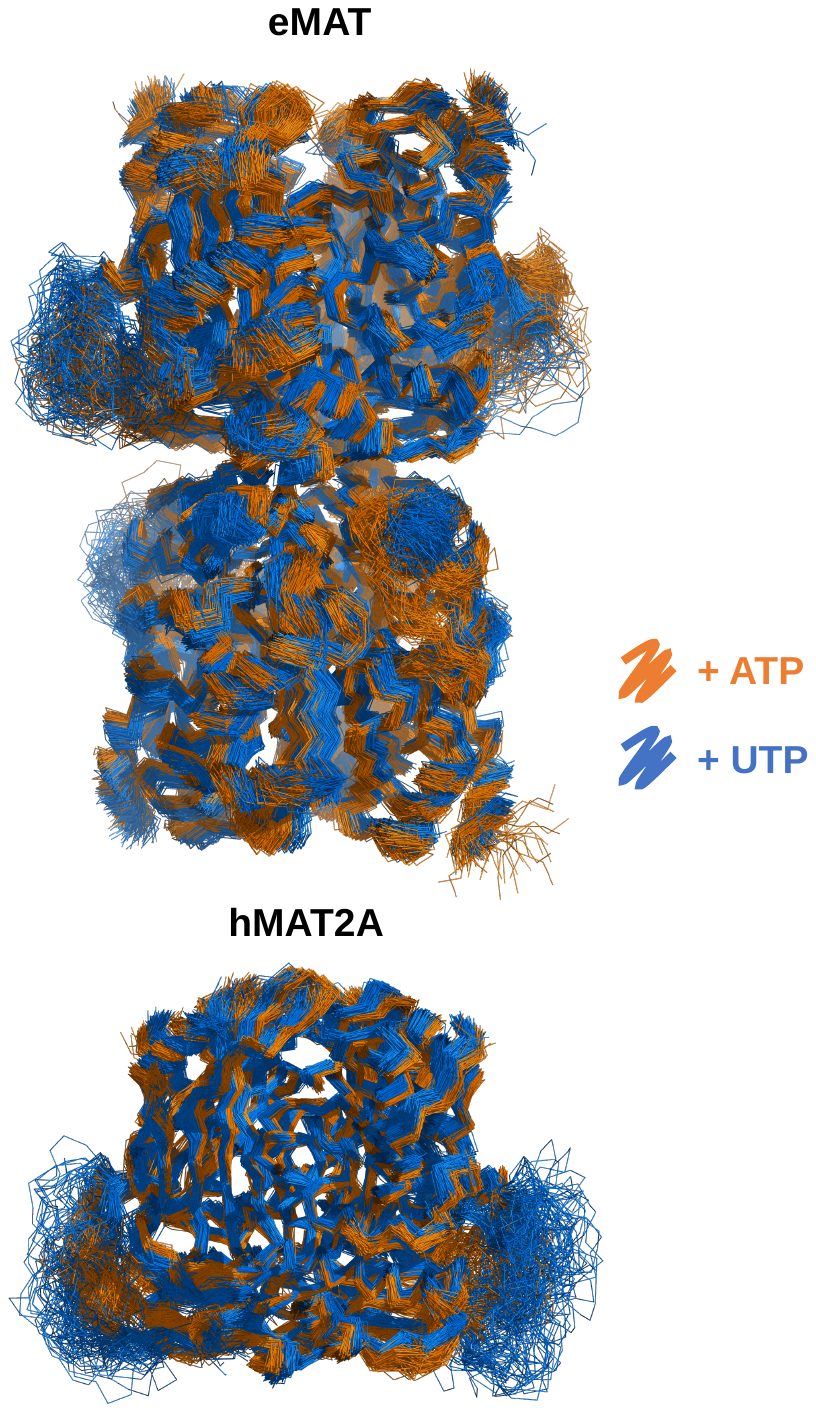

**Supplementary Figure 9. MD backbone ensembles of the eMAT and hMAT2A open states bound to ATP or UTP.** Representative backbone ensembles are shown in a ribbon representation, each consisting of 150 structures sampled evenly over triplicate 500 ns open state simulations. Binding of ATP vs UTP causes no obvious change in conformational sampling throughout either protein backbone except for the active site loop. A broadened conformational sampling of the active site loop in the presence of UTP is observed in both eMAT and hMAT2A.

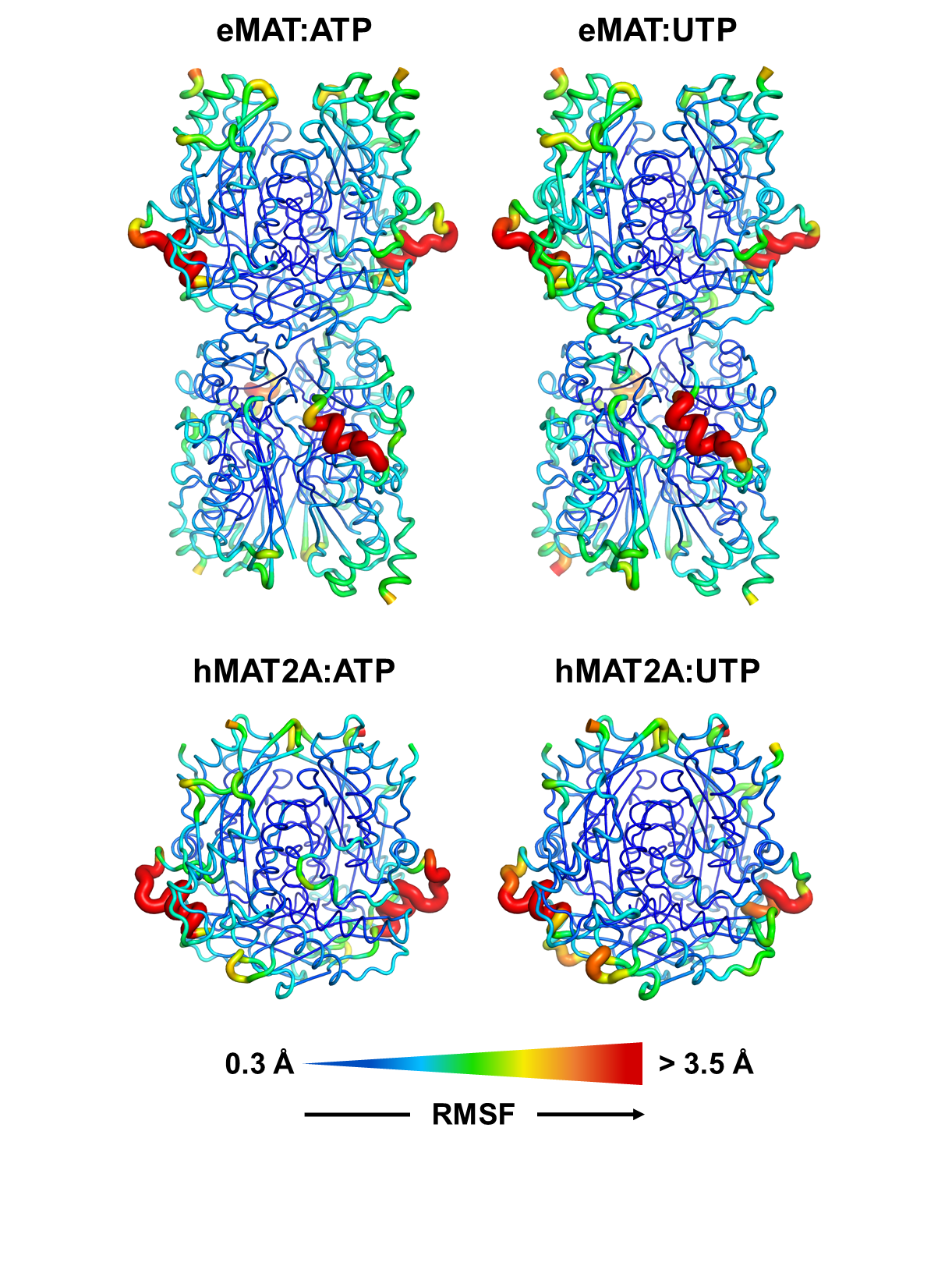

**Supplementary Figure 10. Cα RMSF over open state MD simulation trajectories.** Mapping Cα RMSF values to eMAT and hMAT2A simulation input structures (closed state) highlights the high flexibility observed in the active site loop open state. The RMSF values used were obtained as the average of triplicate trajectories. No obvious differences in eMAT and hMAT2A backbone dynamics were observed over the course of these trajectories.

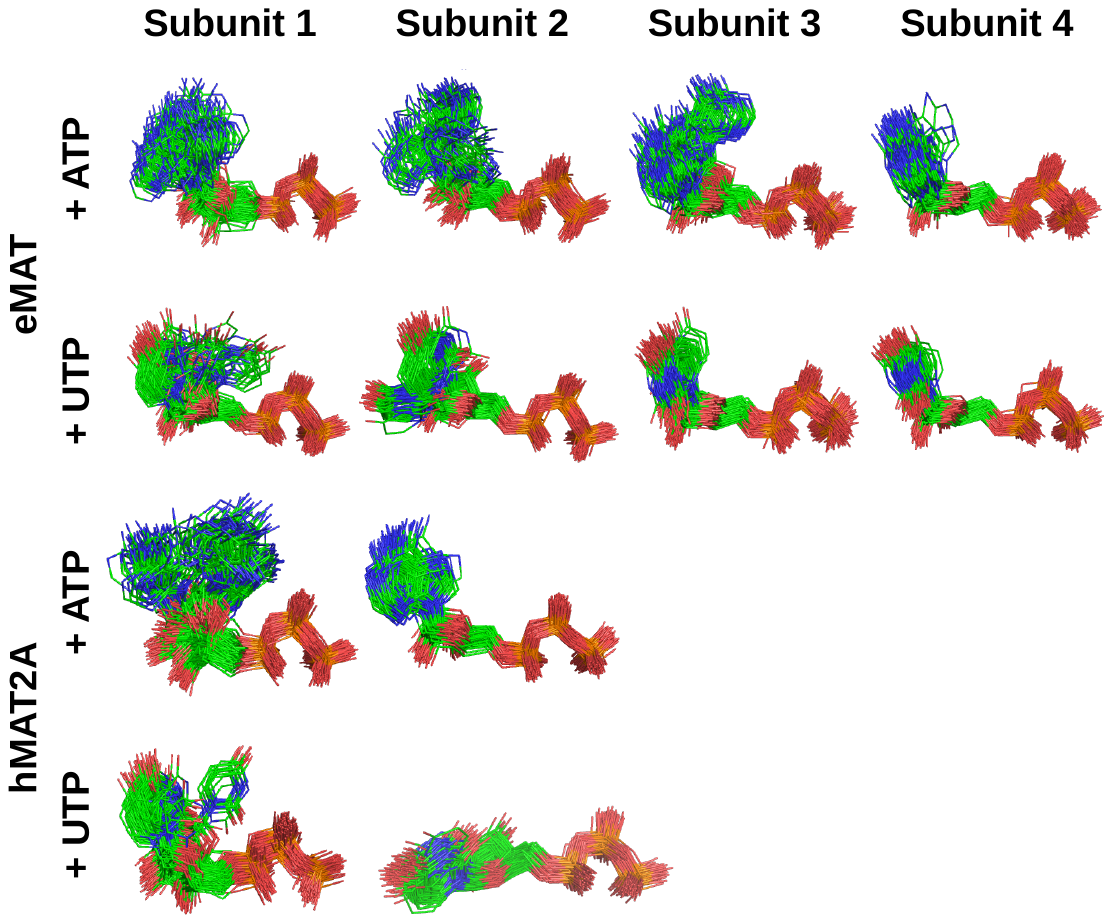

**Supplementary Figure 11. MD substrate ensembles of ATP or UTP bound to eMAT or hMAT2A.** Representative substrate ensembles are shown, each consisting of 150 structures sampled evenly over triplicate 500 ns open state simulations. A protein backbone alignment was used to demonstrate substrate flexibility and conformational variability in the context of the enzyme active site pocket. As the multimeric eMAT and hMAT2A models possess four and two independent active sites, respectively, the substrate poses from each subunit are represented separately.

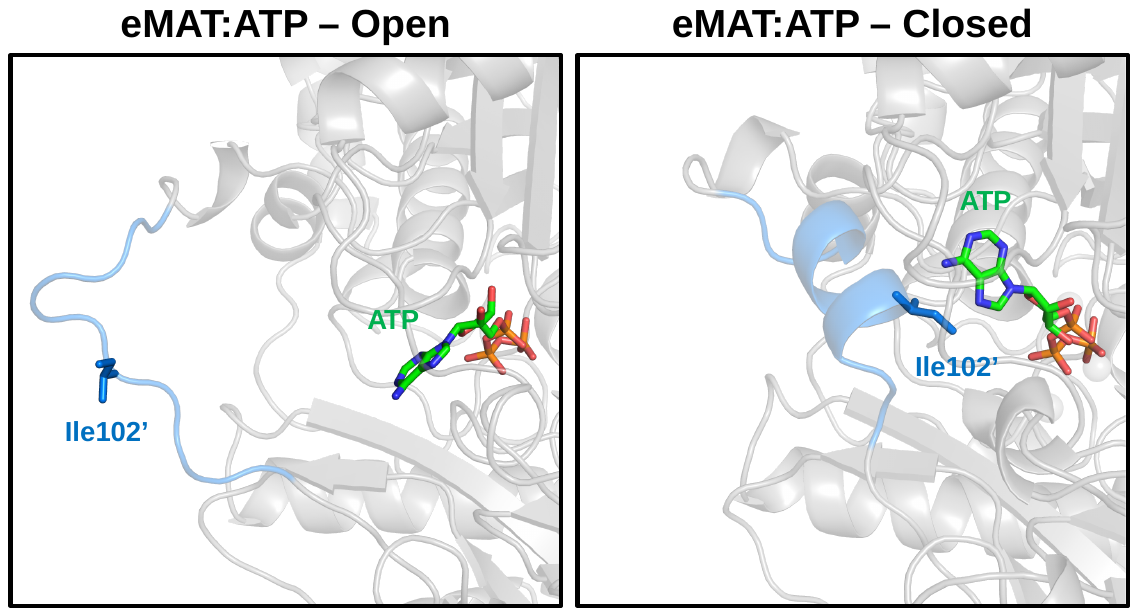

**Supplementary Figure 12. Position of Ile102’ in the open and closed active site loop conformations.** The position of Ile102’ in eMAT:ATP is shown relative to the nucleotide binding site in an arbitrarily selected open state frame and the closed state input structure. In the closed state of the active site loop, this residue is positioned to interact with the nucleobase. In the open state, however, Ile102’ is positioned considerably further away from the nucleobase, preventing stabilizing interactions from occurring.

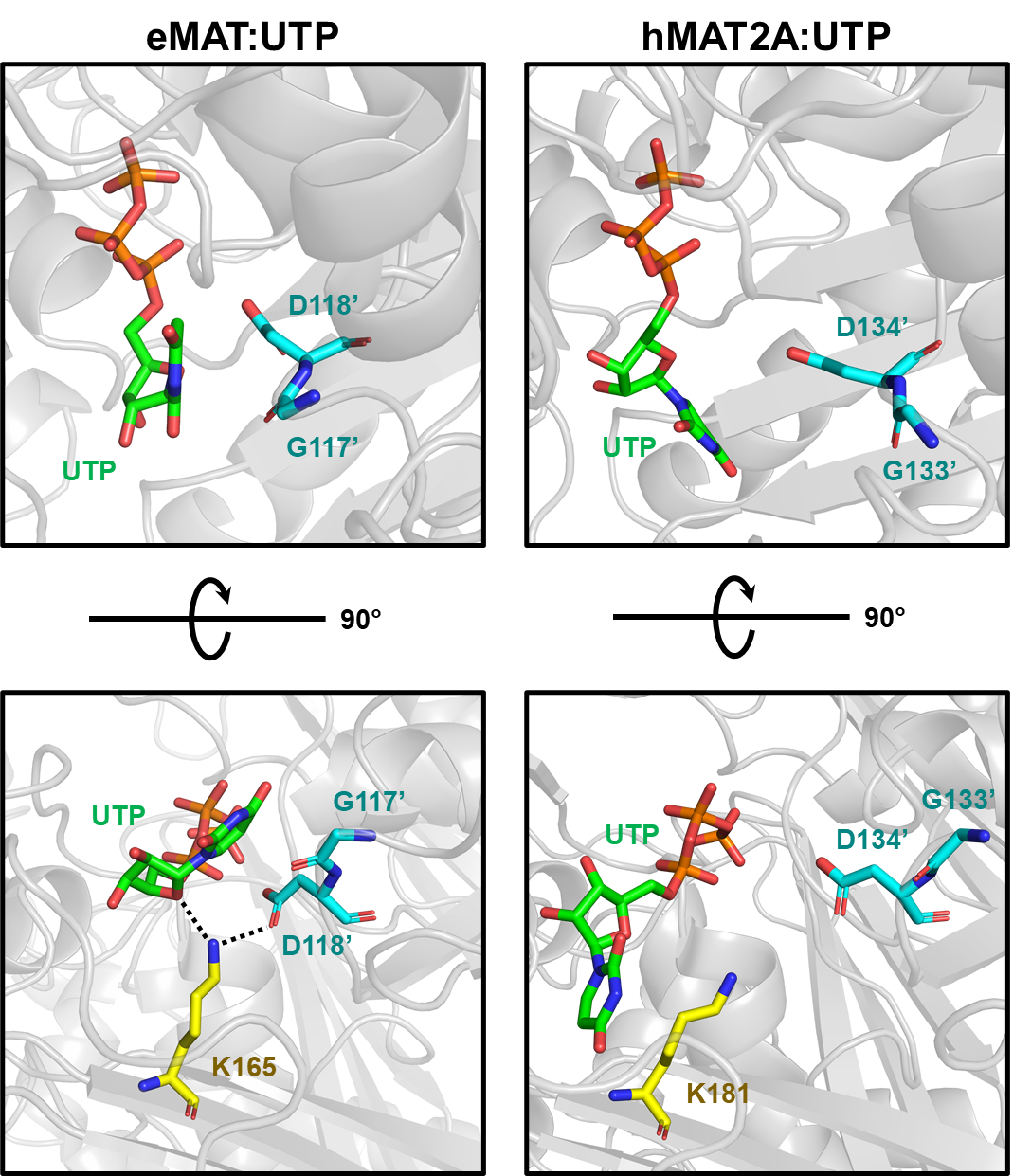

**Supplementary Figure 13. Enzyme-substrate interactions constrain nucleotide conformations**. Electrostatic interactions between the substrate and first shell enzyme residues dictate the accessible UTP conformational space. In the eMAT open state, the close proximity of Gly117’ and Asp118’ to the pyrimidine ring force a strained ꭕ dihedral eclipsed conformation that is not observed in hMAT2A (Figure 4). This conformation is the result of stabilizing interactions between Lys165, Asp118’, and the UTP O4’ and O5’, shown as dotted lines. In the hMAT2A open state, these residues are positioned further from UTP, allowing for the adoption of more relaxed nucleotide conformations.

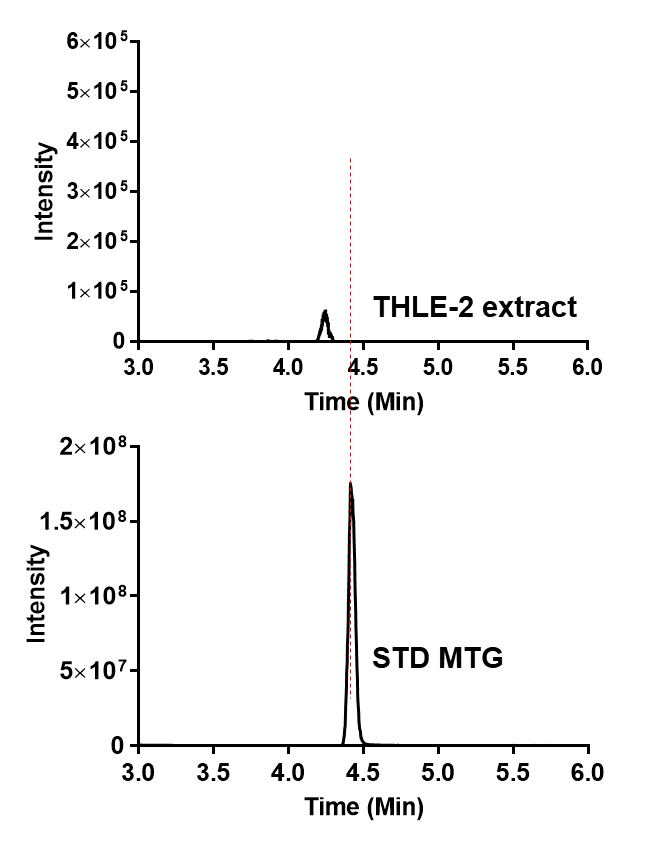

**Supplementary Figure 14.** **LC-MS analysis of metabolite from THLE-2.** Extracted chromatograms of the standard MTG, THLE-2 cell extract. THLE-2 cell extract shows absence of MTG. Experiment was performed in biological triplicate.

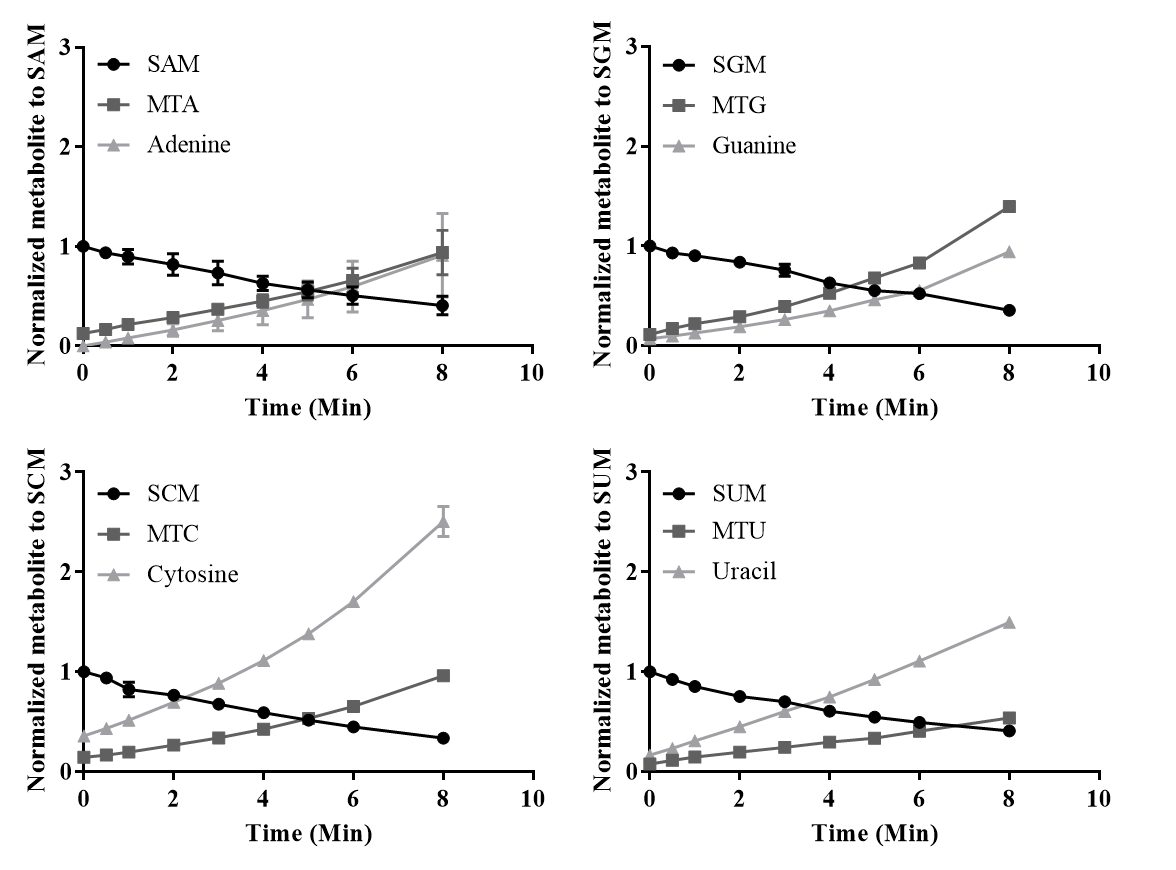

**Supplementary Figure 15. SNM degradation**. SNM analogs were incubated at 37 ̊C and aliquots were taken every hr. Stability of each sample was tested by UPLC. SNM analog degrades to the corresponding MTN and nucleotide bases with respect to the time. Normalized graph with corresponding SNM. Degradation experiments were run in duplicates and error bars show standard deviation.

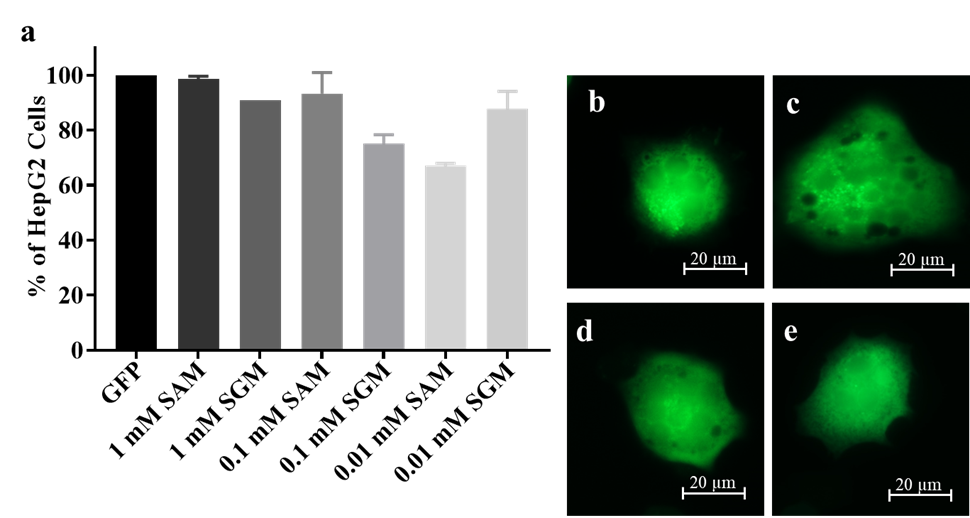

**Supplementary Figure 16. Electroporation of HepG2 cells.** (a) No change in growth rate was observed when HpeG2 cells electroporated with SGM and SAM with pmaxGFP plasmid. Percentage of cells is normalized to cells electroporated with only pmaxGFP plasmid. HepG2 cells electroporated with pmaxGFP plasmid and with (b) 0.1 mM SGM (c) 0.01 mM SGM (d) 0.1 SAM (e) 0.01 mM SAM.

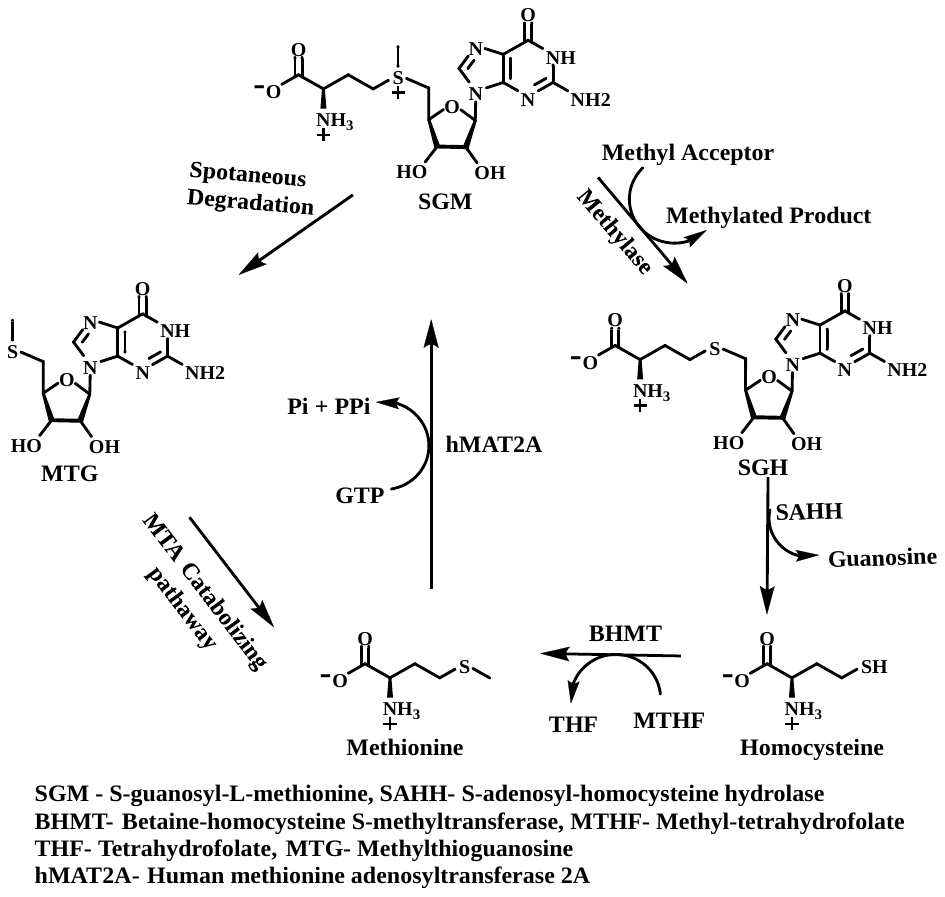

**Supplementary Figure 17. SGM synthesis and possible downstream pathways.** Human methionine adenosyltransferase 2A (hMAT2A) synthesizing S-guanosyl-methionine (SGM) using methionine and GTP. Methyltransferase utilizing SGM as a methyl source to produce methylated product and side product S-guanosyl-homocysteine (SGH). SGH is hydrolyzed by S-adenosyl-homocysteine hydrolase (SAHH) to give homocysteine and guanine. Homocysteine further methylated by methyltetrahydrofolate using betaine homocysteine S-methyltransferase to give methionine. SGH can be spontaneously degraded into the methyl thioguanine (MTG) which is further catabolize into the methionine.

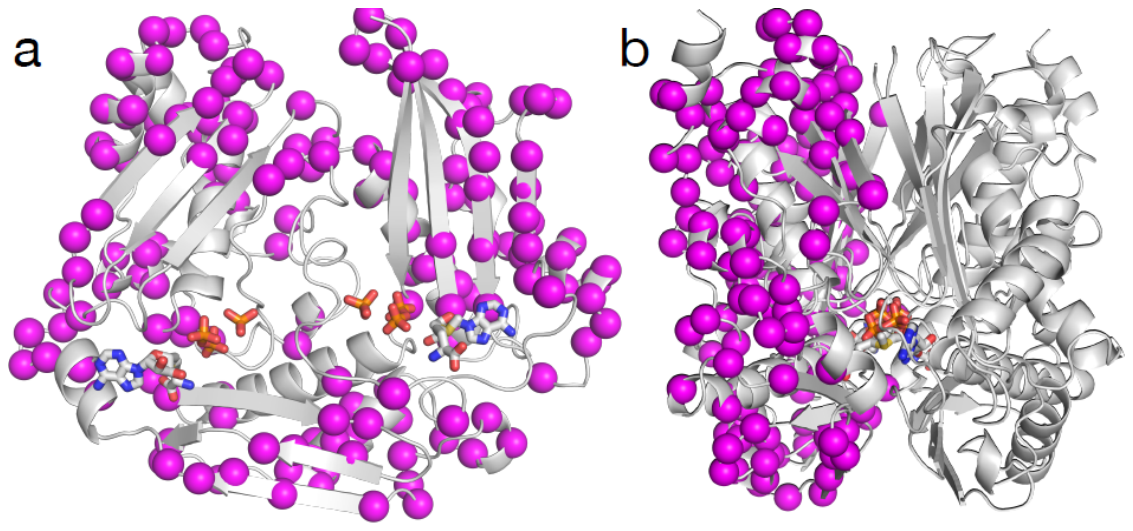

**Supplementary Figure 18. The locations of sequence differences between eMAT and hMAT2A.** The sequence differences are shown in only one monomer of the dimer for clarity. The residues different in the outer shells (i.e., not substrate binding site) are highlighted as purple spheres. The structure of eMAT is shown in grey cartoon representation and products (SAM, PPi, Pi) are shown as sticks. (a) View of the interface of the monomer (“top” monomer in the dimer is omitted for clarity). (b) Lateral view of the dimer.

**Table 1.** **Data collection and refinement statistics**

|  | eMAT:  ATP | eMAT:  GTP | eMAT:  CTP | eMAT:  UTP | eMAT:  UppNHp | eMAT:  GppNHp | hMAT2A:  UppNHp |
| --- | --- | --- | --- | --- | --- | --- | --- |
| **Data collection** |  |  |  |  |  |  |  |
| Space group | P2_1_ | P 6_1_22 | P 2_1_2_1_2_1_ | P 6_1_ 22 | P 6_1_ 22 | P 6_1_ 22 | P 6_3_ 22 |
| Cell dimensions |  |  |  |  |  |  |  |
| *a*, *b*, *c* (Å) | 67.92, 117.78, 113.21 | 122.95, 122.95, 288.09 | 100.14, 118.32, 144.46 | 122.60, 122.60, 288.83 | 123.65, 123.65, 289.41 | 122.34, 122.34, 287.27 | 144.70, 144.70, 187.69 |
| α,β,γ (°) | 90.00, 107.52, 90.00 | 90.00, 90.00, 120.00 | 90.00, 90.00, 90.00 | 90.00, 90.00, 120.00 | 90.00, 90.00, 120.00 | 90.00, 90.00, 120.00 | 90.00, 90.00, 120.00 |
| Resolution (Å) | 39.79-1.95 (1.98-1.95) | 39.86-2.39 (2.46-2.39) | 39.44 - 1.89 (1.92-1.89) | 39.75-2.25 (2.31-2.25) | 43.98-2.24 (2.30-2.24) | 43.65-2.50 (2.59-2.50) | 39.37-2.55 (2.66-2.55) |
| *R*_merge_ | 0.151 (2.148) | 0.258 (5.319) | 0.084 (2.114) | 0.150 (4.079) | 0.114 (6.060) | 0.160 (4.254) | 0.182 (7.635) |
| *R*_pim_ | 0.061 (0.856) | 0.009 (0.851) | 0.033 (0.891) | 0.028 (0.757) | 0.019 (1.025) | 0.026 (0.675) | 0.035 (1.537) |
| *I* / σ*I*  CC(1/2) | 8.2 (1.1)  0.998 (0.445) | 15.5 (1.0)  0.999 (0.434) | 13.8 (0.8)  0.999 (0.412) | 14.2 (1.1)  0.998 (0.549) | 19.6 (0.7)  1.000 (0.385) | 16.2 (1.1)  0.999 (0.622) | 10.4 (0.4)  0.999 (0.571) |
| Completeness (%) | 99.9 (99.8) | 100.0 (100) | 99.9 (99.8) | 100.0 (100) | 100 (100) | 100 (100) | 99.8 (99.7) |
| Multiplicity | 7.0 (7.2) | 39.4 (39.8) | 7.5 (6.3) | 30.3 (30.5) | 38.0 (35.5) | 39.1 (40.3) | 27.0 (25.0) |
| **Refinement** |  |  |  |  |  |  |  |
| Resolution (Å) | 35.99-1.95 (2.02-1.95) | 38.39-2.39 (2.475-2.39) | 38.22-1.89 (1.958-1.89) | 39.09-2.25 (2.33-2.25) | 43.98-2.24 (2.32-2.24) | 43.63-2.5 (2.59-2.5) | 39.40-2.50 (2.56-2.50) |
| Unique reflections | 123330 (12236) | 51784 (5068) | 137216 (13509) | 61629 (6043) | 63616 (6212) | 44820 (4383) | 40352 (2445) |
| *R*_work_ / *R*_free_ | 17.02/20.66 (30.90/32.29) | 18.22/21.57 (30.47/32.88) | 16.59/19.51 (31.49/35.29) | 18.96/21.08 (29.30/33/18) | 21.24/23.46 (34.44/38.08) | 22.29/24.35 (24.35/40.13) | 21.94/24.59 (48.57/52.77) |
| No. atoms | 12593 | 6030 | 12542 | 6140 | 6069 | 6023 | 6146 |
| Protein | 11854 | 5831 | 11671 | 5921 | 5859 | 5894 | 6064 |
| Ligand/ion | 104 | 66 | 148 | 62 | 77 | 49 | 54 |
| Water | 635 | 133 | 633 | 157 | 133 | 80 | 28 |
| *B*-factors |  |  |  |  |  |  |  |
| Protein | 61.77 | 67.34 | 46.05 | 70.46 | 97.25 | 95.26 | 103.91 |
| Ligand/ion | 55.53 | 81.00 | 51.54 | 85.01 | 135.89 | 115.98 | 112.22 |
| Water | 51.78 | 61.54 | 46.50 | 65.37 | 77.40 | 74.58 | 97.28 |
| R.m.s. deviations |  |  |  |  |  |  |  |
| Bond lengths (Å) | 0.011 | 0.003 | 0.010 | 0.006 | 0.004 | 0.002 | 0.002 |
| Bond angles (°) | 0.98 | 0.61 | 1.04 | 0.80 | 0.60 | 0.58 | 0.57 |
| PDB ID | 7LOO | 7LOW | 7LO2 | 7LOZ | 7LL3 | 7LNN | 7LNH |

*All data from single crystals. *Values in parentheses are for highest-resolution shell.

**Supplementary Table 2. Concentrations of NTPs in human^6^ normal and cancer cells, E. coli (mid log phase)^7^.**

| NTP | Normal human cells (mM) | Cancer human cells (mM) | E. coli (mM) |
| --- | --- | --- | --- |
| ATP | 2.537 | 3.134 | 3.560 |
| GTP | 0.232 | 0.473 | 1.660 |
| CTP | 0.083 | 0.402 | 0.325 |
| UTP | 0.227 | 0.686 | 0.667 |

**Supplementary Table 3. The Primers used for mutagenesis of Ser247Thr hMAT2A and Thr227Ser eMAT mutant.**

| Name | Froward Primer^#^ | Reverse Primer |
| --- | --- | --- |
| Ser247Thr hMAT2A mutant | 5'-CAGCCG**ACC**GGTCGTTTCGTTATCGG-3' | 5'-CAGATGATAGATGGTGTCCTCGTCGAG-3' |
| Thr227Ser eMAT mutant | CTTCATCAACCCG**TCT**GGTCGTTTCGTTA | AATTTGGTGGCAGAAGTCAGCCA |

^#^Bold underlined codon in forward primer used for changing Ser to Thr and Thr to Ser.

**hMAT2A plasmid sequence**

ttaatacgactcactataggggaattgtgagcggataacaattcccctctagaaataattttgtttaactttaagaaggagatataccatgggcagcagccatcatcatcatcatcacagcagcggcctggtgccgcgcggcagccatATGAATGGCCAACTGAATGGTTTTCACGAAGCGTTCATCGAAGAGGGCACGTTCCTGTTCACCTCCGAATCTGTTGGTGAAGGTCACCCAGATAAAATCTGCGACCAAATCTCTGACGCGGTGCTGGACGCGCATCTGCAACAGGACCCAGACGCGAAGGTTGCGTGTGAAACCGTCGCTAAGACTGGCATGATCCTGCTGGCTGGCGAAATCACCTCTCGTGCGGCGGTTGACTACCAGAAAGTTGTGCGTGAAGCCGTTAAGCACATCGGCTACGACGACTCTTCTAAAGGTTTCGACTACAAAACCTGTAACGTTCTCGTAGCGCTGGAACAGCAGTCTCCGGACATCGCGCAGGGTGTCCACCTGGACCGTAACGAGGAAGACATCGGTGCGGGCGATCAGGGCCTGATGTTCGGTTATGCGACCGACGAGACTGAGGAATGCATGCCGCTGACCATCGTTCTGGCGCACAAACTCAATGCGAAACTGGCGGAACTGCGTCGTAACGGTACCCTCCCGTGGCTGCGCCCTGACTCTAAAACCCAGGTTACCGTTCAGTACATGCAGGACCGTGGCGCTGTTCTCCCGATCCGTGTCCATACTATCGTGATCTCTGTTCAGCACGATGAAGAAGTTTGCCTGGACGAAATGCGTGACGCCCTCAAAGAAAAAGTTATCAAAGCGGTAGTCCCGGCGAAGTACCTCGACGAGGACACCATCTATCATCTGCAGCCGTCTGGTCGTTTCGTTATCGGTGGTCCGCAAGGCGACGCGGGTCTGACGGGTCGTAAAATCATTGTTGACACCTACGGTGGTTGGGGTGCGCATGGTGGCGGTGCGTTCTCCGGTAAAGACTACACCAAAGTTGACCGTTCCGCGGCATACGCAGCGCGTTGGGTTGCGAAGTCTCTGGTTAAAGGTGGTCTGTGCCGTCGTGTTCTGGTTCAGGTTTCTTACGCAATCGGTGTTTCTCACCCTCTGTCTATCTCTATCTTCCACTATGGTACCTCTCAGAAATCTGAACGTGAGCTGCTGGAAATCGTTAAGAAGAACTTCGACCTGCGTCCGGGTGTGATTGTACGTGACCTGGATCTGAAAAAACCGATCTACCAGCGTACGGCGGCTTACGGCCACTTCGGCCGTGACTCTTTTCCGTGGGAAGTGCCGAAAAAGCTCAAATACtaagaattcgagctccgtcgacaagcttgcggccgcactcgagcaccaccaccaccaccactgagatccggctgctaacaaagcccgaaaggaagctgagttggctgctgccaccgctgagcaataactagcataaccccttggggcctctaaacgggtcttgaggggttttttgctgaaaggaggaactatatccggattggcgaatgggacgcgccctgtagcggcgcattaagcgcggcgggtgtggtggttacgcgcagcgtgaccgctacacttgccagcgccctagcgcccgctcctttcgctttcttcccttcctttctcgccacgttcgccggctttccccgtcaagctctaaatcgggggctccctttagggttccgatttagtgctttacggcacctcgaccccaaaaaacttgattagggtgatggttcacgtagtgggccatcgccctgatagacggtttttcgccctttgacgttggagtccacgttctttaatagtggactcttgttccaaactggaacaacactcaaccctatctcggtctattcttttgatttataagggattttgccgatttcggcctattggttaaaaaatgagctgatttaacaaaaatttaacgcgaattttaacaaaatattaacgtttacaatttcaggtggcacttttcggggaaatgtgcgcggaacccctatttgtttatttttctaaatacattcaaatatgtatccgctcatgaattaattcttagaaaaactcatcgagcatcaaatgaaactgcaatttattcatatcaggattatcaataccatatttttgaaaaagccgtttctgtaatgaaggagaaaactcaccgaggcagttccataggatggcaagatcctggtatcggtctgcgattccgactcgtccaacatcaatacaacctattaatttcccctcgtcaaaaataaggttatcaagtgagaaatcaccatgagtgacgactgaatccggtgagaatggcaaaagtttatgcatttctttccagacttgttcaacaggccagccattacgctcgtcatcaaaatcactcgcatcaaccaaaccgttattcattcgtgattgcgcctgagcgagacgaaatacgcgatcgctgttaaaaggacaattacaaacaggaatcgaatgcaaccggcgcaggaacactgccagcgcatcaacaatattttcacctgaatcaggatattcttctaatacctggaatgctgttttcccggggatcgcagtggtgagtaaccatgcatcatcaggagtacggataaaatgcttgatggtcggaagaggcataaattccgtcagccagtttagtctgaccatctcatctgtaacatcattggcaacgctacctttgccatgtttcagaaacaactctggcgcatcgggcttcccatacaatcgatagattgtcgcacctgattgcccgacattatcgcgagcccatttatacccatataaatcagcatccatgttggaatttaatcgcggcctagagcaagacgtttcccgttgaatatggctcataacaccccttgtattactgtttatgtaagcagacagttttattgttcatgaccaaaatcccttaacgtgagttttcgttccactgagcgtcagaccccgtagaaaagatcaaaggatcttcttgagatcctttttttctgcgcgtaatctgctgcttgcaaacaaaaaaaccaccgctaccagcggtggtttgtttgccggatcaagagctaccaactctttttccgaaggtaactggcttcagcagagcgcagataccaaatactgtccttctagtgtagccgtagttaggccaccacttcaagaactctgtagcaccgcctacatacctcgctctgctaatcctgttaccagtggctgctgccagtggcgataagtcgtgtcttaccgggttggactcaagacgatagttaccggataaggcgcagcggtcgggctgaacggggggttcgtgcacacagcccagcttggagcgaacgacctacaccgaactgagatacctacagcgtgagctatgagaaagcgccacgcttcccgaagggagaaaggcggacaggtatccggtaagcggcagggtcggaacaggagagcgcacgagggagcttccagggggaaacgcctggtatctttatagtcctgtcgggtttcgccacctctgacttgagcgtcgatttttgtgatgctcgtcaggggggcggagcctatggaaaaacgccagcaacgcggcctttttacggttcctggccttttgctggccttttgctcacatgttctttcctgcgttatcccctgattctgtggataaccgtattaccgcctttgagtgagctgataccgctcgccgcagccgaacgaccgagcgcagcgagtcagtgagcgaggaagcggaagagcgcctgatgcggtattttctccttacgcatctgtgcggtatttcacaccgcatatatggtgcactctcagtacaatctgctctgatgccgcatagttaagccagtatacactccgctatcgctacgtgactgggtcatggctgcgccccgacacccgccaacacccgctgacgcgccctgacgggcttgtctgctcccggcatccgcttacagacaagctgtgaccgtctccgggagctgcatgtgtcagaggttttcaccgtcatcaccgaaacgcgcgaggcagctgcggtaaagctcatcagcgtggtcgtgaagcgattcacagatgtctgcctgttcatccgcgtccagctcgttgagtttctccagaagcgttaatgtctggcttctgataaagcgggccatgttaagggcggttttttcctgtttggtcactgatgcctccgtgtaagggggatttctgttcatgggggtaatgataccgatgaaacgagagaggatgctcacgatacgggttactgatgatgaacatgcccggttactggaacgttgtgagggtaaacaactggcggtatggatgcggcgggaccagagaaaaatcactcagggtcaatgccagcgcttcgttaatacagatgtaggtgttccacagggtagccagcagcatcctgcgatgcagatccggaacataatggtgcagggcgctgacttccgcgtttccagactttacgaaacacggaaaccgaagaccattcatgttgttgctcaggtcgcagacgttttgcagcagcagtcgcttcacgttcgctcgcgtatcggtgattcattctgctaaccagtaaggcaaccccgccagcctagccgggtcctcaacgacaggagcacgatcatgcgcacccgtggggccgccatgccggcgataatggcctgcttctcgccgaaacgtttggtggcgggaccagtgacgaaggcttgagcgagggcgtgcaagattccgaataccgcaagcgacaggccgatcatcgtcgcgctccagcgaaagcggtcctcgccgaaaatgacccagagcgctgccggcacctgtcctacgagttgcatgataaagaagacagtcataagtgcggcgacgatagtcatgccccgcgcccaccggaaggagctgactgggttgaaggctctcaagggcatcggtcgagatcccggtgcctaatgagtgagctaacttacattaattgcgttgcgctcactgcccgctttccagtcgggaaacctgtcgtgccagctgcattaatgaatcggccaacgcgcggggagaggcggtttgcgtattgggcgccagggtggtttttcttttcaccagtgagacgggcaacagctgattgcccttcaccgcctggccctgagagagttgcagcaagcggtccacgctggtttgccccagcaggcgaaaatcctgtttgatggtggttaacggcgggatataacatgagctgtcttcggtatcgtcgtatcccactaccgagatatccgcaccaacgcgcagcccggactcggtaatggcgcgcattgcgcccagcgccatctgatcgttggcaaccagcatcgcagtgggaacgatgccctcattcagcatttgcatggtttgttgaaaaccggacatggcactccagtcgccttcccgttccgctatcggctgaatttgattgcgagtgagatatttatgccagccagccagacgcagacgcgccgagacagaacttaatgggcccgctaacagcgcgatttgctggtgacccaatgcgaccagatgctccacgcccagtcgcgtaccgtcttcatgggagaaaataatactgttgatgggtgtctggtcagagacatcaagaaataacgccggaacattagtgcaggcagcttccacagcaatggcatcctggtcatccagcggatagttaatgatcagcccactgacgcgttgcgcgagaagattgtgcaccgccgctttacaggcttcgacgccgcttcgttctaccatcgacaccaccacgctggcacccagttgatcggcgcgagatttaatcgccgcgacaatttgcgacggcgcgtgcagggccagactggaggtggcaacgccaatcagcaacgactgtttgcccgccagttgttgtgccacgcggttgggaatgtaattcagctccgccatcgccgcttccactttttcccgcgttttcgcagaaacgtggctggcctggttcaccacgcgggaaacggtctgataagagacaccggcatactctgcgacatcgtataacgttactggtttcacattcaccaccctgaattgactctcttccgggcgctatcatgccataccgcgaaaggttttgcgccattcgatggtgtccgggatctcgacgctctcccttatgcgactcctgcattaggaagcagcccagtagtaggttgaggccgttgagcaccgccgccgcaaggaatggtgcatgcaaggagatggcgcccaacagtcccccggccacggggcctgccaccatacccacgccgaaacaagcgctcatgagcccgaagtggcgagcccgatcttccccatcggtgatgtcggcgatataggcgccagcaaccgcacctgtggcgccggtgatgccggccacgatgcgtccggcgtagaggatcgagatctcgatcccgcgaaa

**eMAT plasmid sequence**

ttaatacgactcactataggggaattgtgagcggataacaattcccctctagaaataattttgtttaactttaagaaggagatataccatgggcagcagccatcatcatcatcatcacagcagcggcctggtgccgcgcggcagccatATGGCAAAACACCTTTTTACGTCCGAGTCCGTCTCTGAAGGGCATCCTGACAAAATTGCTGACCAAATTTCTGATGCCGTTTTAGACGCGATCCTCGAACAGGATCCGAAAGCACGCGTTGCTTGCGAAACCTACGTAAAAACCGGCATGGTTTTAGTTGGCGGCGAAATCACCACCAGCGCCTGGGTAGACATCGAAGAGATCACCCGTAACACCGTTCGCGAAATTGGCTATGTGCATTCCGACATGGGCTTTGACGCTAACTCCTGTGCGGTTCTGAGCGCTATCGGCAAACAGTCTCCTGACATCAACCAGGGCGTTGACCGTGCCGATCCGCTGGAACAGGGCGCGGGTGACCAGGGTCTGATGTTTGGCTACGCAACTAATGAAACCGACGTGCTGATGCCAGCACCTATCACCTATGCACACCGTCTGGTACAGCGTCAGGCTGAAGTGCGTAAAAACGGCACTCTGCCGTGGCTGCGCCCGGACGCGAAAAGCCAGGTGACTTTTCAGTATGACGACGGCAAAATCGTTGGTATCGATGCTGTCGTGCTTTCCACTCAGCACTCTGAAGAGATCGACCAGAAATCGCTGCAAGAAGCGGTAATGGAAGAGATCATCAAGCCAATTCTGCCCGCTGAATGGCTGACTTCTGCCACCAAATTCTTCATCAACCCGACCGGTCGTTTCGTTATCGGTGGCCCAATGGGTGACTGCGGTCTGACTGGTCGTAAAATTATCGTTGATACCTACGGCGGCATGGCGCGTCACGGTGGCGGTGCATTCTCTGGTAAAGATCCATCAAAAGTGGACCGTTCCGCAGCCTACGCAGCACGTTATGTCGCGAAAAACATCGTTGCTGCTGGCCTGGCCGATCGTTGTGAAATTCAGGTTTCCTACGCAATCGGCGTGGCTGAACCGACCTCCATCATGGTAGAAACTTTCGGTACTGAGAAAGTGCCTTCTGAACAACTGACCCTGCTGGTACGTGAGTTCTTCGACCTGCGCCCATACGGTCTGATTCAGATGCTGGATCTGCTGCACCCGATCTACAAAGAAACCGCAGCATACGGTCACTTTGGTCGTGAACATTTCCCGTGGGAAAAAACCGACAAAGCGCAGCTGCTGCGCGATGCTGCCGGTCTGAAGTAActcgagcaccaccaccaccaccactgagatccggctgctaacaaagcccgaaaggaagctgagttggctgctgccaccgctgagcaataactagcataaccccttggggcctctaaacgggtcttgaggggttttttgctgaaaggaggaactatatccggattggcgaatgggacgcgccctgtagcggcgcattaagcgcggcgggtgtggtggttacgcgcagcgtgaccgctacacttgccagcgccctagcgcccgctcctttcgctttcttcccttcctttctcgccacgttcgccggctttccccgtcaagctctaaatcgggggctccctttagggttccgatttagtgctttacggcacctcgaccccaaaaaacttgattagggtgatggttcacgtagtgggccatcgccctgatagacggtttttcgccctttgacgttggagtccacgttctttaatagtggactcttgttccaaactggaacaacactcaaccctatctcggtctattcttttgatttataagggattttgccgatttcggcctattggttaaaaaatgagctgatttaacaaaaatttaacgcgaattttaacaaaatattaacgtttacaatttcaggtggcacttttcggggaaatgtgcgcggaacccctatttgtttatttttctaaatacattcaaatatgtatccgctcatgaattaattcttagaaaaactcatcgagcatcaaatgaaactgcaatttattcatatcaggattatcaataccatatttttgaaaaagccgtttctgtaatgaaggagaaaactcaccgaggcagttccataggatggcaagatcctggtatcggtctgcgattccgactcgtccaacatcaatacaacctattaatttcccctcgtcaaaaataaggttatcaagtgagaaatcaccatgagtgacgactgaatccggtgagaatggcaaaagtttatgcatttctttccagacttgttcaacaggccagccattacgctcgtcatcaaaatcactcgcatcaaccaaaccgttattcattcgtgattgcgcctgagcgagacgaaatacgcgatcgctgttaaaaggacaattacaaacaggaatcgaatgcaaccggcgcaggaacactgccagcgcatcaacaatattttcacctgaatcaggatattcttctaatacctggaatgctgttttcccggggatcgcagtggtgagtaaccatgcatcatcaggagtacggataaaatgcttgatggtcggaagaggcataaattccgtcagccagtttagtctgaccatctcatctgtaacatcattggcaacgctacctttgccatgtttcagaaacaactctggcgcatcgggcttcccatacaatcgatagattgtcgcacctgattgcccgacattatcgcgagcccatttatacccatataaatcagcatccatgttggaatttaatcgcggcctagagcaagacgtttcccgttgaatatggctcataacaccccttgtattactgtttatgtaagcagacagttttattgttcatgaccaaaatcccttaacgtgagttttcgttccactgagcgtcagaccccgtagaaaagatcaaaggatcttcttgagatcctttttttctgcgcgtaatctgctgcttgcaaacaaaaaaaccaccgctaccagcggtggtttgtttgccggatcaagagctaccaactctttttccgaaggtaactggcttcagcagagcgcagataccaaatactgtccttctagtgtagccgtagttaggccaccacttcaagaactctgtagcaccgcctacatacctcgctctgctaatcctgttaccagtggctgctgccagtggcgataagtcgtgtcttaccgggttggactcaagacgatagttaccggataaggcgcagcggtcgggctgaacggggggttcgtgcacacagcccagcttggagcgaacgacctacaccgaactgagatacctacagcgtgagctatgagaaagcgccacgcttcccgaagggagaaaggcggacaggtatccggtaagcggcagggtcggaacaggagagcgcacgagggagcttccagggggaaacgcctggtatctttatagtcctgtcgggtttcgccacctctgacttgagcgtcgatttttgtgatgctcgtcaggggggcggagcctatggaaaaacgccagcaacgcggcctttttacggttcctggccttttgctggccttttgctcacatgttctttcctgcgttatcccctgattctgtggataaccgtattaccgcctttgagtgagctgataccgctcgccgcagccgaacgaccgagcgcagcgagtcagtgagcgaggaagcggaagagcgcctgatgcggtattttctccttacgcatctgtgcggtatttcacaccgcatatatggtgcactctcagtacaatctgctctgatgccgcatagttaagccagtatacactccgctatcgctacgtgactgggtcatggctgcgccccgacacccgccaacacccgctgacgcgccctgacgggcttgtctgctcccggcatccgcttacagacaagctgtgaccgtctccgggagctgcatgtgtcagaggttttcaccgtcatcaccgaaacgcgcgaggcagctgcggtaaagctcatcagcgtggtcgtgaagcgattcacagatgtctgcctgttcatccgcgtccagctcgttgagtttctccagaagcgttaatgtctggcttctgataaagcgggccatgttaagggcggttttttcctgtttggtcactgatgcctccgtgtaagggggatttctgttcatgggggtaatgataccgatgaaacgagagaggatgctcacgatacgggttactgatgatgaacatgcccggttactggaacgttgtgagggtaaacaactggcggtatggatgcggcgggaccagagaaaaatcactcagggtcaatgccagcgcttcgttaatacagatgtaggtgttccacagggtagccagcagcatcctgcgatgcagatccggaacataatggtgcagggcgctgacttccgcgtttccagactttacgaaacacggaaaccgaagaccattcatgttgttgctcaggtcgcagacgttttgcagcagcagtcgcttcacgttcgctcgcgtatcggtgattcattctgctaaccagtaaggcaaccccgccagcctagccgggtcctcaacgacaggagcacgatcatgcgcacccgtggggccgccatgccggcgataatggcctgcttctcgccgaaacgtttggtggcgggaccagtgacgaaggcttgagcgagggcgtgcaagattccgaataccgcaagcgacaggccgatcatcgtcgcgctccagcgaaagcggtcctcgccgaaaatgacccagagcgctgccggcacctgtcctacgagttgcatgataaagaagacagtcataagtgcggcgacgatagtcatgccccgcgcccaccggaaggagctgactgggttgaaggctctcaagggcatcggtcgagatcccggtgcctaatgagtgagctaacttacattaattgcgttgcgctcactgcccgctttccagtcgggaaacctgtcgtgccagctgcattaatgaatcggccaacgcgcggggagaggcggtttgcgtattgggcgccagggtggtttttcttttcaccagtgagacgggcaacagctgattgcccttcaccgcctggccctgagagagttgcagcaagcggtccacgctggtttgccccagcaggcgaaaatcctgtttgatggtggttaacggcgggatataacatgagctgtcttcggtatcgtcgtatcccactaccgagatatccgcaccaacgcgcagcccggactcggtaatggcgcgcattgcgcccagcgccatctgatcgttggcaaccagcatcgcagtgggaacgatgccctcattcagcatttgcatggtttgttgaaaaccggacatggcactccagtcgccttcccgttccgctatcggctgaatttgattgcgagtgagatatttatgccagccagccagacgcagacgcgccgagacagaacttaatgggcccgctaacagcgcgatttgctggtgacccaatgcgaccagatgctccacgcccagtcgcgtaccgtcttcatgggagaaaataatactgttgatgggtgtctggtcagagacatcaagaaataacgccggaacattagtgcaggcagcttccacagcaatggcatcctggtcatccagcggatagttaatgatcagcccactgacgcgttgcgcgagaagattgtgcaccgccgctttacaggcttcgacgccgcttcgttctaccatcgacaccaccacgctggcacccagttgatcggcgcgagatttaatcgccgcgacaatttgcgacggcgcgtgcagggccagactggaggtggcaacgccaatcagcaacgactgtttgcccgccagttgttgtgccacgcggttgggaatgtaattcagctccgccatcgccgcttccactttttcccgcgttttcgcagaaacgtggctggcctggttcaccacgcgggaaacggtctgataagagacaccggcatactctgcgacatcgtataacgttactggtttcacattcaccaccctgaattgactctcttccgggcgctatcatgccataccgcgaaaggttttgcgccattcgatggtgtccgggatctcgacgctctcccttatgcgactcctgcattaggaagcagcccagtagtaggttgaggccgttgagcaccgccgccgcaaggaatggtgcatgcaaggagatggcgcccaacagtcccccggccacggggcctgccaccatacccacgccgaaacaagcgctcatgagcccgaagtggcgagcccgatcttccccatcggtgatgtcggcgatataggcgccagcaaccgcacctgtggcgccggtgatgccggccacgatgcgtccggcgtagaggatcgagatctcgatcccgcgaaa
